# Lead (Pb) exposure alters neural cell fate in the developing human brain

**DOI:** 10.1101/2025.08.14.670354

**Authors:** Maureen M. Sampson, Sayli J. Sonsurkar, Lisa Nieland, Lin Pan, Ryan L. Kan, Daria J. Azizad, Xiang Li, Nardos Kebede, Alicia R. Lane, Priya E. D’Souza, Parinya Panuwet, Emily J. Hill, Samantha N. Lanjewar, Anna N. Voss, Dorelle V. Fawwal, Anson Sing, Alexia King, Erica Werner, Jennifer M. Spangle, Dana Boyd Barr, Aparna Bhaduri, Ye Zhang, Victor Faundez, Steven A. Sloan

## Abstract

The heavy metal lead (Pb) is a developmental neurotoxicant associated with cognitive and behavioral deficits, but the cellular mechanisms underlying these impairments remain unclear. Here we show that prenatal Pb exposure biases human radial glia fate, prolonging neurogenesis and suppressing astrogenesis. We used hiPSC-derived cortical organoids, primary human fetal tissue, and *in vivo* xenografts to demonstrate that Pb exposure alters radial glial differentiation. Pb-exposed organoids contain a higher proportion of neurons and fewer astrocytes. We validated this differentiation bias in primary radial glia from human cortices (GW16-20), observing Pb-associated reductions in astrocyte commitment via genetic lineage tracing. This correlated with increased H3K27me3, a repressive histone modification deposited by the histone methyltransferase complex PRC2, suggesting epigenetic reprogramming as a mechanistic link between Pb and neural cell fate commitment. Our findings indicate that prenatal Pb exposure impacts lineage commitment in the developing brain, which may contribute to cognitive and behavioral impairment.

## Introduction

The brain is most vulnerable during the prenatal period of rapid expansion, when both genetic and environmental perturbations can disrupt when and where differentiation occurs. In the prenatal period, neural progenitor cells called radial glia sequentially differentiate into three major cell classes: neurons, astrocytes, and oligodendrocytes. In the human cortex, radial glia begin neurogenesis by the 6^th^ gestational week (GW6), giving rise to both neuronally committed intermediate progenitors and post-mitotic neurons^1,2^. By gestational week (GW) 15, radial glia cell fate shifts towards glial progenitors, astrocytes, and oligodendrocytes in a process called the gliogenic switch^3–5^. Factors that regulate radial glia differentiation are critical orchestrators of brain development because they determine the timing and quantity of cells that eventually comprise the mature brain. For this reason, abnormal progenitor proliferation and differentiation are implicated in many neurodevelopmental disorders such as intellectual disability and autism spectrum disorder^6,7^.

One of the most well established developmental neurotoxicants is the heavy metal lead (Pb). The molecular mechanisms of Pb toxicity have been studied extensively in rodents and human cell lines^8–10^; however, the cellular phenotypes that contribute to behavioral and cognitive delays in humans remain unknown. In the US, approximately 2.5% of pregnant women exhibit high blood Pb levels exceeding 2 µg/dL^11^, highlighting Pb exposure as an ongoing public health concern. Prenatal exposure to Pb is associated with neonatal behavioral and cognitive deficits^9,10^. For example, elevated maternal Pb blood levels during the 2^nd^ trimester are associated with an increased risk for cognitive developmental delay^12^. Furthermore, maternal blood Pb levels during pregnancy are more predictive of 2-year neurodevelopmental scores in offspring than postnatal measurements in the offspring^13^, suggesting the critical period for Pb exposure may be prenatal.

In mice, chronic perinatal Pb exposure increased oligodendrocyte precursor cells, suggesting that Pb can alter progenitor stemness and cell fate^14^. Previous work using human neural progenitor cells characterized Pb-induced changes in gene expression, epigenetics, and cell fate^15–17^. In directed cultures of embryonic stem cell-derived neural progenitor cells, Pb exposure induced premature neurogenesis, ultimately increasing the generation of neurons.^15^ Understanding how Pb toxicity leads to neural progenitor dysfunction is critical because it affects all progenies including neurons, astrocytes, oligodendrocytes, ependymal cells, and other cell types of neuroepithelial origin.

Modeling cell fate transitions of multipotent cells like radial glia are affected is challenging in 2D culture because their differentiation trajectories change from neurogenesis to gliogenesis throughout development. To address this, our study leveraged a more sophisticated *in vitro* approach: 3D hiPSC cortical organoids, which better reflect the complexity and expansive nature of radial glia subytpes^18,19^. Organoids can also be maintained over long time periods (months to years), and this permits spontaneous generation of radial glia subtypes that encompass both neurogenic^20^ and gliogenic^19,21^ cell fate decisions. This is particularly relevant to human neurodevelopmental studies because human brain development occurs over longer a longer temporal scale than rodent models or 2D cultures can be maintained. Lastly, the use of human iPSCs allows for the possibility to study human-specific gene-environment interactions.

As a reductionist model system, organoids are best suited to test specific, hypothesis-driven toxicodynamic questions. They are especially useful when findings can be validated with primary human tissue samples. Initial studies using organoids to assess Pb-dependent cell fate changes have suggested disruption of neuronal commitment and maturation^22,23^. Here, we used cortical organoids to first examine shifting progenitor fate commitment during Pb exposure. We chose to focus our study on the cortex, which performs high level cognitive function. With regard to the timing of Pb exposure in cortical neural progenitors^24^ we targeted a developmental timepoint in corticogenesis that encompasses the generation of two major cell types: excitatory neurons and astrocytes. We modeled a mid-second trimester exposure to capture peak neurogenesis and early gliogenesis (fetal GW16-18 ≈ organoid D130-D150)^18,19,21,25^. Organoids can be maintained over longer timeframes than 2D cultures and therefore better reflect real-world exposure durations; however, it remains critical to establish biologically relevant dosing paradigms. In our study we first carefully optimized exposure conditions to reflect meaningful levels observed in mammalian exposures by benchmarking against previously reported brain tissue Pb burdens. This involved extensive titration and validation to ensure that Pb dosing in organoids mimicked real-world exposure scenarios, enabling meaningful comparisons across models.

Overall, the approach to validate our exposure paradigm, both dosing and exposure period, can be applied to many other exposures and serves as a framework for many future studies in conjunction with a few pioneering studies have used organoids to examine neurodevelopmental toxicants^26–28^. The hiPSC organoid approach continues to advance, but currently only primary tissues include the full cellular diversity of the developing human brain, and some subtypes of radial glia are absent from organoids^29,30^ and rodents^31,32^. We therefore validated our findings using orthogonal approaches in primary human fetal brain, as well as xenografts of Pb-exposed fetal radial glia into developing mice. Together with previous work^26^, our study further demonstrates the applicability of brain organoids for studying the molecular mechanisms of developmental toxicants.

## Results

### Primary human and organoid neural cells accumulate intracellular Pb at similar rates

Human *in vitro* models are valuable tools for neurotoxicology due to limited access to human brain tissue at relevant developmental stages. In this work we modeled Pb toxicity in hiPSC cortical organoids^19,25,33^, which recapitulate many aspects of cortical development including astrogenesis (Extended Data Fig. 1a-b). To evaluate organoids as a translatable model for Pb toxicodynamic studies, we first asked whether neural lineage cells within cortical organoids accumulate Pb at similar rates to primary fetal human populations. We purified astrocytes and neurons from both organoids (D150-D175) and primary fetal cortex (GW18-19) by immunopanning using anti-HepaCAM and anti-CD24 antibodies respectively. We then visualized uptake of Pb-acetate in 2D cultures of neurons or astrocytes using live imaging and the fluorescent heavy metal sensor leadmium^34^ (Fig. 1a). Both fetal and organoid populations rapidly accumulated Pb within minutes and intracellular Pb levels stabilized in less than an hour (Fig. 1b, n=8 cells, see Extended Data Fig. 1c for individual traces). We observed no significant differences in Pb uptake between organoid and fetal astrocytes or neurons (Fig. 1c, Brown-Forsythe ANOVA test and Dunnetts multiple comparison test). Control additions of saline or acetate (vehicle control) did not elicit leadmium fluorescence greater than 1 DF/F, consistent with baseline drift, whereas PbA-low (500 nM) and PbA-high (500 µM) averaged 3.0 and 28.9 DF/F respectively (Extended Data Fig. 1d).

**Fig. 1.**
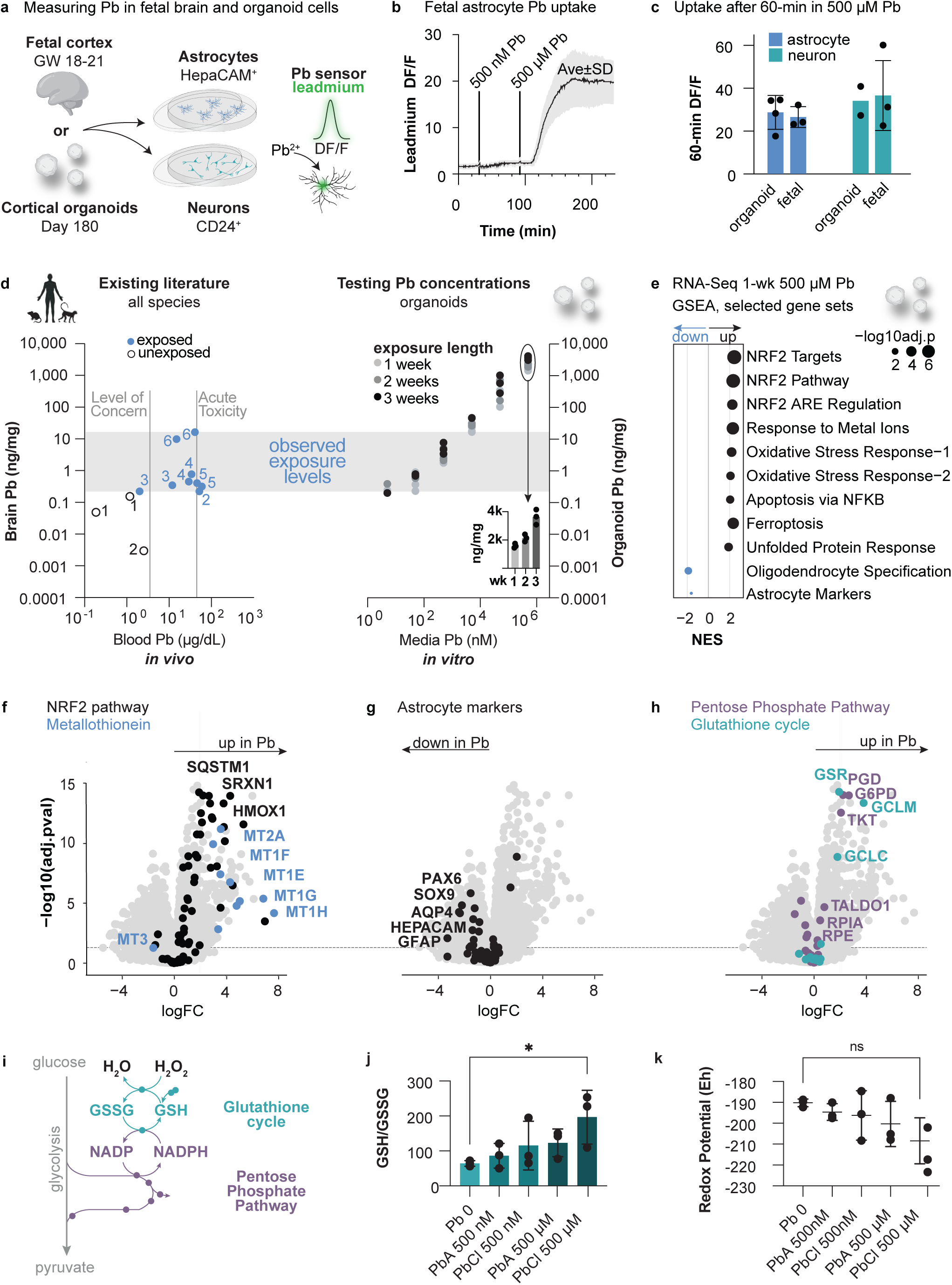
Cortical organoids provide a platform for studying developmental neurotoxicity of the heavy metal lead (Pb). **a**, Fetal and organoid-derived astrocytes and neurons were isolated by immunopanning using anti-HepaCAM and anti-CD24 antibodies, respectively. The fluorescent lead sensor leadmium was used to measure Pb uptake in purified cells. **b**, A representative leadmium trace from GW18 fetal astrocytes exposed to PbA-low (500 nM) then PbA-high (500 µM). Data is plotted as baseline-normalized leadmium fluorescence (DF/F) using a 20 min baseline. **c**, DF/F after 60-min exposure to Pb for fetal and organoid-derived cells. Each data point is the average of ∼8 cells from a single organoid differentiation or tissue sample (N=2 hiPSC lines and 3 organoid differentiations and N=3 fetal tissues). Error bars are mean and SD, statistical test is Brown-Forsythe ANOVA test and Dunnetts multiple comparison test. **d**, Brain tissue Pb-burdens from references^35–40^, respectively, are compared to cortical organoid Pb levels after exposure to Pb for 1-3 weeks. Each data point is a single literature value (left) or tissue measurement from a single organoid (right). (**d**, inset) Pb measurements in organoid tissue show dose- and duration-dependent Pb increases even at maximal PbA-high (500 µM) exposures. **e**, GSEA analysis using Hallmark pathways for cortical organoids exposed to PbCl_2_ for 1 week. **f-h**, Volcano plots highlighting specific differentially expressed genes related to (**f**) NRF2 pathway and metallothionines, (**g**) astrocyte markers, (**h**) glutathione cycle and pentose phosphate pathway. **i**, A schematic of the glutathione cycle and pentose phosphate pathway. **j-k**, Reduced glutathione (GSH) and oxidized (GSSG) glutathione was measured in whole organoid lysate by mass spectrometry. Each data point represents a single analytical sample from 3 pooled organoids. Antioxidant status is shown as the ratio of GSH/GSSG in **j** and the calculated redox potential in **k**. Error bars are mean and SD for all panels. The statistical comparison in (**j**) is Dunnett’s multiple comparisons test following a 1-way ANOVA.

### Organoids accumulate toxicologically relevant Pb tissue burdens that induce antioxidant response after 1 week

A critical consideration in toxicology research is the dose and delivery of the exposure. Therefore, we carefully consulted the literature to define a human/mammal exposure range of brain parenchyma Pb levels to inform our *in vitro* exposure strategy. Our approach was to first define a tissue-level Pb target from animal/human studies, then empirically determine which Pb exposure levels *in vitro* result in similar organoid tissue Pb accumulation. There is limited data on Pb burdens in brain tissue because exposures are almost exclusively reported as blood Pb levels. Therefore, to put published brain tissue Pb burdens in context, we plotted tissue burdens with paired blood measurements in Fig. 1d. Many of these measurements were reported from controlled exposures for toxicology studies in non-human primates^35,36^ and rodents^37–39^, but we also included a study that used human post-mortem tissues^40^. Brain tissues from Pb exposed animals ranged from 0.2-16.5 ng/mg (indicated as grey bar in Fig. 1d) and their paired blood measurements ranged between the blood lead level of concern as defined by the CDC^41^ (3.5 µg/dL = 50 ppb) and acutely toxic levels of lead (40 µg/dL).

Next, we proceeded to establish our own dosing strategy titrated to achieve organoid tissue Pb burdens that recapitulated the literature-informed toxicologically relevant ranges in Fig. 1d. We grew organoids in media with concentrations of Pb acetate spanning 6 orders of magnitude (5 nM – 500 µM) and quantified tissue Pb burdens using inductively coupled plasma mass spectrometry (ICP-MS) (Fig. 1d). For all Pb doses, tissue measurements increased linearly over the 3-week exposures, including the highest exposure group of 500 µM Pb (Fig. 1d right panel, inset), suggesting cellular capacity for continual Pb accumulation even at very high Pb levels. Organoids exposed to 500 nM Pb acetate resulted in organoid tissue Pb levels most closely approximating *in vivo* brain tissue Pb burdens, ranging from 1.79-7.8 ng/mg (average 2.4-5.3 ng/mg) across the 3 week-exposure (Fig. 1d). For this reason, we perform exposure experiments with “Pb-low” 500 nM Pb (103.6 ppb or 10.36 ug/dL) to recapitulate biological *in vivo* exposures or with “Pb-high” 500 µM Pb to examine maximal Pb effects.

Inorganic Pb is available as a salt with acetate or halide groups, and previous toxicology studies have almost exclusively used the acetate salt due to its higher solubility in aqueous solutions. To examine the impact of different Pb salts, we performed RNA-Seq on whole organoids (D157-D163) grown in control media or media containing Pb acetate (PbA) or PbCl_2_ for 1 week. Overall, PbCl_2_ induced a stronger transcriptomic response than PbA (Extended Data Fig. 1e). Gene set enrichment analysis (GSEA) with hallmark pathways showed consistent overrepresentation of genes associated with reactive oxygen species, MTORC1, apoptosis, TNFA signaling via NFKB, xenobiotic metabolism, inflammation, and oxidative phosphorylation in the setting of Pb exposure (Extended Data Fig. 1f). Broadly, Pb-high exposure (500 µM) induced a strong transcriptional response in organoids, but Pb-low (500 nM) had more subtle effects. GSEA with curated pathways (gene set C2) showed classical markers of toxicity including cell death and oxidative stress but also revealed strongly activated neuroprotective programs including the NRF2 antioxidant pathway, metal response proteins, and unfolded protein response (Fig.1e). Pb exposure increased all metallothionein genes except MT3 (Fig.1f), which is highly expressed in astrocytes^42,43^. Interestingly, upon Pb exposure, we observed a decrease in both astrocyte markers and oligodendrocyte specification genes including canonical markers SOX9, AQP4, HEPACAM, GFAP, OLIG1 and OLIG2 (Fig.1g).

The transcription factor NRF2 coordinates the antioxidant response to oxidative stress. In the presence of Pb, we observed a consistent increase in genes related to the antioxidant glutathione and the pentose phosphate pathway, which is required to restore reducing equivalents of glutathione (Fig. 1h-i). To test whether glutathione levels and redox status were dysregulated in the presence of Pb, we measured reduced (GSH) and oxidized (GSSG) glutathione in whole organoid lysate. The ratio of GSH/GSSG was increased in PbA-high (500 µM) organoids (P=0.0349 by Dunnett’s multiple comparisons test, N=3 organoids), suggesting a decrease in redox potential (Fig. 1j-k).

### Chronic exposure to PbA-low upregulated genes associated with MTORC1, hypoxia and UPR

One week of Pb-high exposure (500 µM) induced canonical toxicity responses, but most human Pb exposures occur at lower concentrations over longer time frames. Therefore, we next exposed cortical organoids to PbA-low for 3 weeks from D130-D150 (Fig. 2a). To examine cell-specific responses to Pb, we performed droplet-based single-cell sequencing (10X Genomics) on the final day of Pb exposure. A total of 38,562 cells (from N=3 hiPSC lines/experiments) passed QC after sequencing (Fig 2a, Extended Data Fig. 2a-c). We observed all major cell types previously described in the cortical organoid model including excitatory neurons, (few) inhibitory neurons, radial glia, astrocytes, and ependymal cells (Fig. 2b-c). Cell clusters were manually identified based on differential gene expression profiles (Fig. 2d). In differentiated astrocytes and neurons, significantly enriched GSEA hallmark pathways largely overlapped with PbA-high (500 µM) acute exposures in our previous bulk organoid dataset (Fig. 2e and Fig 1e, overlapping pathways marked by *). As an exception, acute exposures to Pb-high (500 µM) upregulated genes associated with increases in oxidative phosphorylation genes, while chronic, Pb-low (500 nM) was associated with an increase in glycolytic genes. “MTORC1 signaling” was the top GSEA hit in both astrocytes and neurons.

**Fig. 2.**
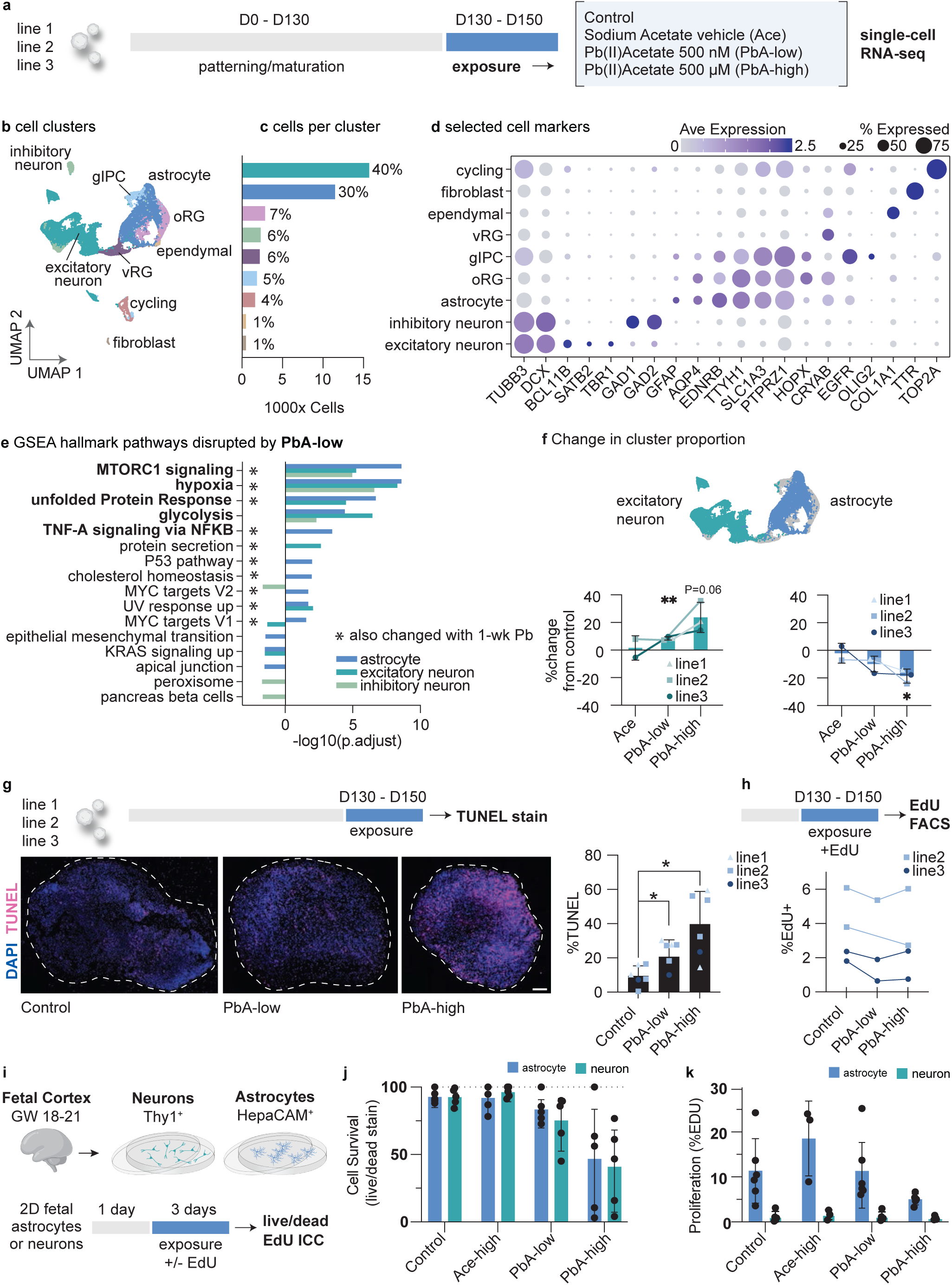
Pb exposure is associated with an increased abundance of excitatory neurons and decrease numbers of astrocytes that is not explained by changes in cell survival or proliferation. **a,** Cortical organoids (N=3 hiPSC lines) were dissociated and single cells captured after 3 weeks of Pb exposure (D130-D150). **b**, A UMAP of 38,562 cells individual cells that passed QC and were assigned to clusters. **c**, The number of cells per cluster and their proportions as a percentage. **d**, Cell identity was determined manually based on differentially expressed genes for each cluster. A subset of selected marker genes used for cell identity are shown. **e**, Hallmark pathways gene-set enrichment analysis for excitatory neurons, inhibitory neurons, and astrocytes. **f**, The percent change in the proportion of excitatory neurons or astrocytes for acetate alone or PbA exposures relative to unexposed control organoids for each hiPSC line. Statistical tests are a one sample t test and wilcoxon test to compare experimental groups to the control “0”. **g**, Cell death in organoids quantified with TUNEL staining after 3 weeks of Pb exposure (D130-D150). Each data point represents the TUNEL staining for an individual organoid (N=3 hiPSC lines with n=1-3 technical replicates per line). Statistical tests were performed including individual replicates: Brown-Forsythe ANOVA test P=0.0096, Dunnett’s multiple comparisons test P=0.013 and P=0.019. **h**, Proliferation in organoids quantified by EdU staining after 3 weeks of EdU + Pb exposure (D130-D150). EdU+ cells were quantified by flow cytometry. Each point represents the percentage of EdU+ cells of total singlet cells (each sample represents 5 pooled organoids). **i**, Fetal neurons (teal) and astrocytes (blue) were immunopanned from cortical brain tissue. Purified cell types were grown in 2D and exposed to Pb or control media for 3 days before live/dead imaging with the calcium indicator calcein and membrane impermeable dye ethidium homodimer to quantify survival. Concurrent cultures were grown with EdU to quantify proliferation. **j**, Neuronal (teal) and astrocyte (blue) survival after 3-day Pb exposure quantified by live/dead imaging. Each point represents a unique tissue sample. **k**, Neuronal (teal) and astrocyte (blue) proliferation in culture over 3-day Pb exposure measured by EdU ICC. Each point represents a unique tissue sample. For **j-k**, comparisons are multiple unpaired t-tests comparing astrocyte vs neuron groups corrected for FDR. All error bars are mean and SD.

### Pb exposure is associated with an increase in excitatory neurons and decrease in astrocytes

Pb exposure altered the percentage of cells assigned to the astrocyte and excitatory neuron clusters in a dose-dependent manner that was consistent across all three hiPSC lines (Fig. 2f). The proportion of cells assigned to the astrocyte cluster decreased as the concentration of Pb increased with -18.6% change for PbA-high (500 µM) (PbA-high vs control, P=0.024, two-tailed t-test). Acetate alone did not have this effect (P=0.75). In contrast, excitatory neuron proportions increased with Pb concentration (P=0.0067 PbA-low (500 nM) vs control, and P=0.065 PbA-high (500 µM) vs control). The Pb-associated decrease in astrocytes is consistent with our previous finding that acute Pb decreased astrocyte gene expression in whole organoid RNA-Seq (Fig. 1e, g).

We next examined whether the Pb associated change in cell lineage proportions was driven by cell type-specific biases in survival or proliferation. We first used approaches to measure gross organoid cell death and proliferation, before proceeding to cell lineage-specific effects. To quantify cell death, we used TUNEL staining, which marks DNA damage. We observed a general Pb-dependent increase in cell death from 23% (PbA-low, 500 nM) and 39.9% (PbA-high, 500 µM) (Fig. 2g, Brown-Forsythe ANOVA test P=0.0096, Dunnett’s multiple comparisons test P=0.013 and P=0.019). To measure proliferation, we exposed organoids to the nucleotide analog EdU throughout the 3-week Pb exposure to identify new-born cells with newly synthesized DNA. We observed no Pb-dependent changes in proliferation in organoids formed from 2 hiPSC lines in 2 differentiations. Instead, cell proliferation rates were largely correlated only with hiPSC line (Fig. 2h).

To address whether there were biases in survival of specific neural lineages, and to ensure we were not examining organoid-specific artifacts, we isolated astrocytes and neurons from primary fetal tissue and measured survival and proliferation after 3 days of Pb exposure (Fig. 2i). We did not observe a difference in survival by cell type (PbA-low 83% vs 75% surival, PbA-high (500 µM) 47% vs 41% survival) using the calcium indicator calcein and membrane impermeable dye ethidium homodimer (Fig. 2j). Using the proliferation marker EdU, we found that astrocytes proliferated in culture and the rate of cell division was unchanged with PbA-low exposure and decreased with PbA-high (500 µM), likely a result of increased cell death (Fig. 2k). As expected, neurons exhibited low levels of proliferation in culture even in the absence of Pb exposure (Fig. 2k). Together, these data suggest our previous observations of diminished astrocyte populations in Pb-exposed organoids are not explained simply by lineage-specific biases in astrocyte proliferation or toxicity.

### Pb exposure modulated putative PRC2 gene targets and increases H3K27me3 deposition

Our organoid Pb exposures reflect second trimester human fetal development, a time window when radial glia are capable of generating both glia and neurons (Fig. 3a). We therefore hypothesized that Pb alters radial glia cell fate decisions in organoids by suppressing astrogenesis. To explore this possibility, we specifically examined Pb-induced transcriptional changes in progenitor cell clusters using both GSEA and gene ontology (GO). All three progenitor populations (oRG, vRG, gIPC) showed overrepresentation of MTORC1 genes (Extended Data Fig. 3a). The oRG cluster upregulated genes in terms related to neuron differentiation, neuron fate commitment, regulation of gliogenesis, cell fate commitment, and Notch signaling pathway (Fig. 3b).

**Fig. 3.**
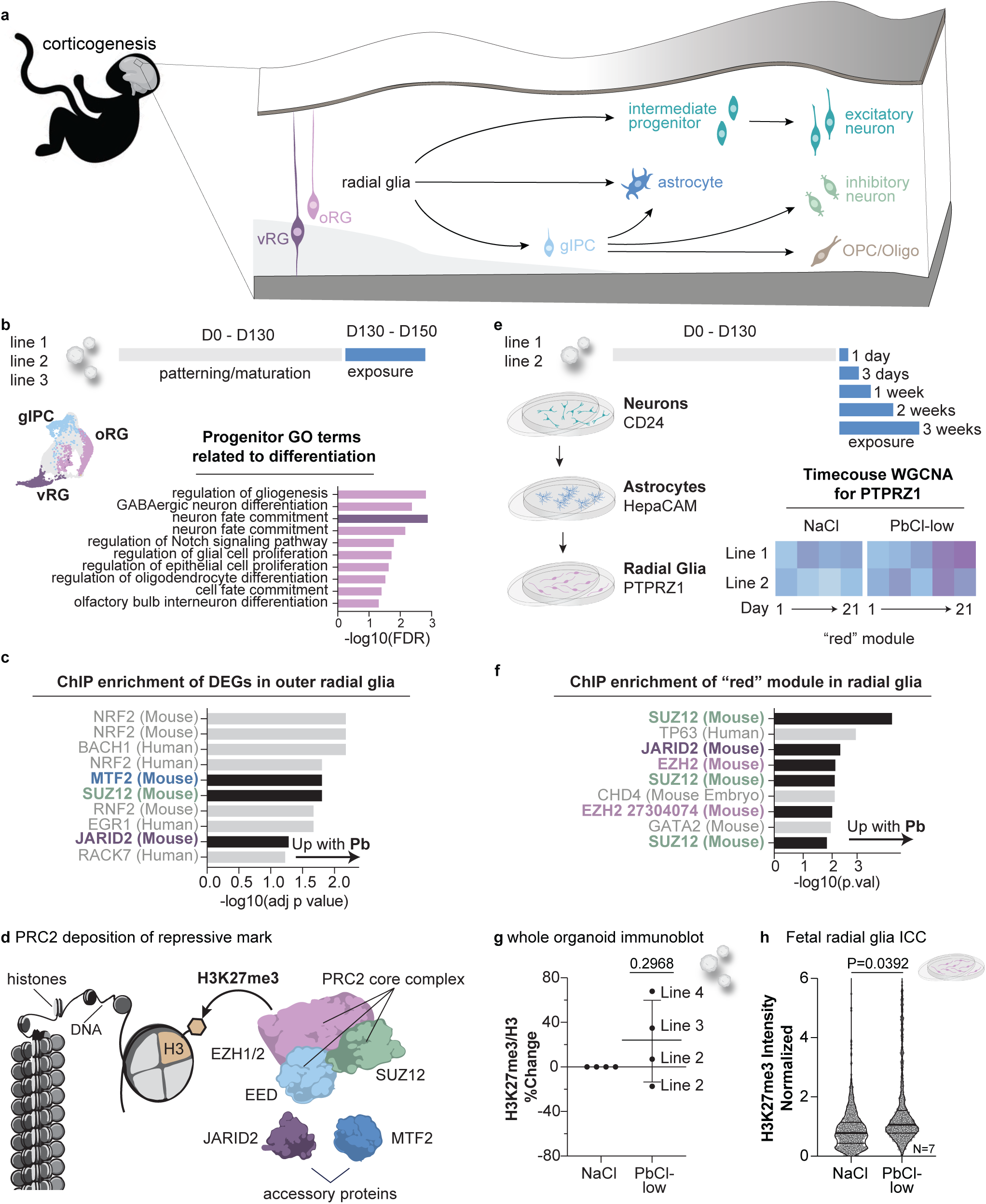
Putative PRC2 gene targets and H3K27me3 increases over Pb exposure. **a**, Radial glia are capable of generating many cell lineages during corticogenesis. **b**, GO terms related to differentiation for single-cell progenitor clusters after 3-week exposure to Pb from D130-D150. **c**, ChIP enrichment analysis (ChEA) of the oRG cluster of cells. **d**, PRC2 deposits repressive the H3K27me3 post translational modification. JARID2 and MTF2 are accessory proteins that recruit the main core complex to DNA targets. **e**, Organoids were exposed to Pb for 3 weeks and immunopanning was performed to isolate radial glia at 5 timepoints for RNA-Seq. WGCNA module-trait relationships for PTPRZ1 immunopanned cells revealed a “red” module of genes that increased over the exposure period. **f**, ChIP enrichment analysis (ChEA) showed an overrepresentation of PRC2-associated genes in the red module. **g**, Immunoblot for H3K27me3/H3 in whole organoid lysate at D150 after 3 weeks of PbCl_2_ exposure, Pb-exposed organoid H3K27me3/H3 is normalized to NaCl. Each point represents a immunoblot from a different organoid differentiation. **h,** PTPRZ1+ radial glia were immunopanned from fetal cortex (GW16-21, N=7 tissue samples), exposed to NaCl or PbCl-low for 3 days, then stained for anti-H3K27me3. The nuclear intensities of H3K27me3 labeling were normalized to the average NaCl nuclei intensity for each sample (N=7). For visualization in **h**, 145 nuclei were subsampled from each sample (N=7) and individual nuclei data are shown in the violin plot, which also shows the median (thick bar) and quartiles. A one-sample t-test was performed for the N=7 average PbCl_2_-low nuclei intensity for each sample (P=0.0392) as shown in Extended Data Fig. 3f).

To investigate transcription factors and epigenetic modifying proteins that have the potential to shift cell fate, we employed ChIP enrichment analysis (ChEA) on the oRG DEGs. ChEA identified NRF2 and BACH1 as the top DNA-binding proteins associated with the DEGs. NRF2 binds a regulatory enhancer called the ‘antioxidant response element,’ which is competitively bound by BACH1 to negatively regulate the antioxidant response. This analysis also identified three components of the Polycomb repressive complex 2 (PRC2): SUZ12, MTF2, and JARID2 (Fig 3c). PRC2 is an epigenetic regulator that deposits repressive H3K27me3, a post translational histone modification that is implicated in neuronal fate^44,45^. SUZ12 is a core component of PRC2, while MTF2 and JARID2 associate with the core PRC2 complex and recruit to target loci (Fig 3d).

We reasoned that the Pb-induced changes we observed in the scRNA-Seq dataset could be biased by overpowered single cell comparisons (despite N = 3 hiPSC lines). To examine how oRGs respond to Pb change over the course of exposure we grew organoids (D130 organoids, N = 2 hiPSC lines) in media containing PbCl_2_-low (500 nM) or NaCl (1 µM)) and subsequently immunopanned neurons, astrocytes, and outer radial glia at 24-hour, 3-day, 1-week, 2-week, and 3-week timepoints (Fig. 3e) for RNA-Seq. Our immunopanning strategy showed enrichment for cell-specific markers as expected (Extended Data Fig. 3b). To identify gene network patterns over the exposure duration we employed weighted gene co-expression network analysis (WGCNA) (Extended Data Fig. 3c). We found one particular module (red) in radial glia (PTPRZ1+ cells) containing genes that exhibited linearly increasing expression with Pb exposure duration (Fig. 3e). ChEA of these genes again identified dysregulation of core PRC2 components. These include SUZ12, EZH2, as well as the PRC2 recruitment protein JARID2 (Fig. 3f), which closely matched the findings from our (independent) single-cell dataset (Fig. 3c).

PRC2 has previously been implicated in the regulation of neurogenesis, astrogenesis and oligogenesis^44–46^.Therefore, we next tested whether H3K27me3, which is deposited by PRC2, was altered by Pb exposure. We exposed organoids from 3 hiPSC lines to PbCl_2_-low for 3 weeks (D130-D150). At D150 we performed acid extraction of histones and measured levels of the transcriptionally repressive mark H3K27me3 by immunoblot. Pb exposed organoids demonstrated increased H3K27me3 levels in 3 of 4 experiments, though this did not reach significance by one-way t test or ANOVA (P=0.2968, Fig. 3g, Extended Data Fig. 3d-e). To examine whether H3K27me3 was changed in neural progenitors specifically, we used the same immunopanning scheme to isolate PTPRZ1 cells from fetal cortical tissue (GW16-21). PTPRZ1 cells were exposed to NaCl or PbCl_2_-low for 3 days then stained with DAPI and anti-H3K27me3 antibody. We quantified H3K27me3 immunocytochemistry signal in individual nuclei (n=18,197 nuclei from N=7 fetal samples) and found that PbCl_2_-low exposed cells had H3K27me3 signals that were 1.78 times more intense than NaCl-exposed cells (one-way t-test, P=0.0392, N=7 normalized PbCl_2_-low averages, see Fig 3h and Extended Data Fig. 3f).

### Primary fetal-derived radial glia generate fewer newborn astrocytes in Pb exposure

To directly test whether radial glia cell fate is altered by Pb, we enriched for radial glia from primary fetal cortices (GW16-21) using immunopanning with either anti-LIFR or anti-PTPRZ1 antibodies after depletion of neurons and astrocytes (Fig 4. a-b). Both immunopanning antibodies increased the percentage of cells expressing the radial glia marker PAX6 (from 7.5% before panning to 34.6% (LIFR) and 25.5% (PTPRZ1) after panning) (Fig. 4b). Cell yields were significantly higher when we immunopanned with PTPRZ1 with an average yield of 19.9 million cells (N=12, PTPRZ1) vs 205 thousand cells (N=12, LIFR) per sample. After 1 day recovery *in vitro*, we exposed radial glia cultures to either PbCl_2_-low, PbA-low, or control media for 3 days. Overall, Pb-low (500 nM) exposures did not change cell proliferation or survival rates beyond sample-to-sample variation (Fig. 4c). After 3 days of differentiation in culture, we immunostained for astrocyte (SOX9) and neuronal cell markers (TUJ1) (Extended Data Fig.4a). We also used EdU to identify new-born cells and quantified SOX9 intensity to identify those committed towards an astrocyte fate (Fig. 4d-e). We observed SOX9 expression in both radial glia and astrocytes, with increased expression correlating with astrocyte lineage commitment. To address this spectrum of expression, we measured SOX9 intensity for individual nuclei, as shown for a representative sample (GW 17) in Fig. 4e. We also defined a threshold for SOX9 positivity and then calculated the percent of EdU+ nuclei that exceeded this fluorescence intensity (Fig. 4e). We found that with Pb exposure 8 out of 10 samples showed a decrease in EdU+SOX9+ cells with an average of -26% loss for PbCl_2_-low and -21% loss for PbA-low (Fig. 4f, one-sample t-test P=0.015 for PbCl_2_-low and P=0.071 for PbA-low). Astrogenic fate was most consistently suppressed at GW16-18 and the effect was more attenuated at GW20-21 for PbCl_2_-exposed cells (Extended Data Fig.4b). We also observed the same decrease in astrocyte fate acquisition irrespective of our radial glia immunopanning antibody choice (Extended Data Fig.4c).

**Fig. 4.**
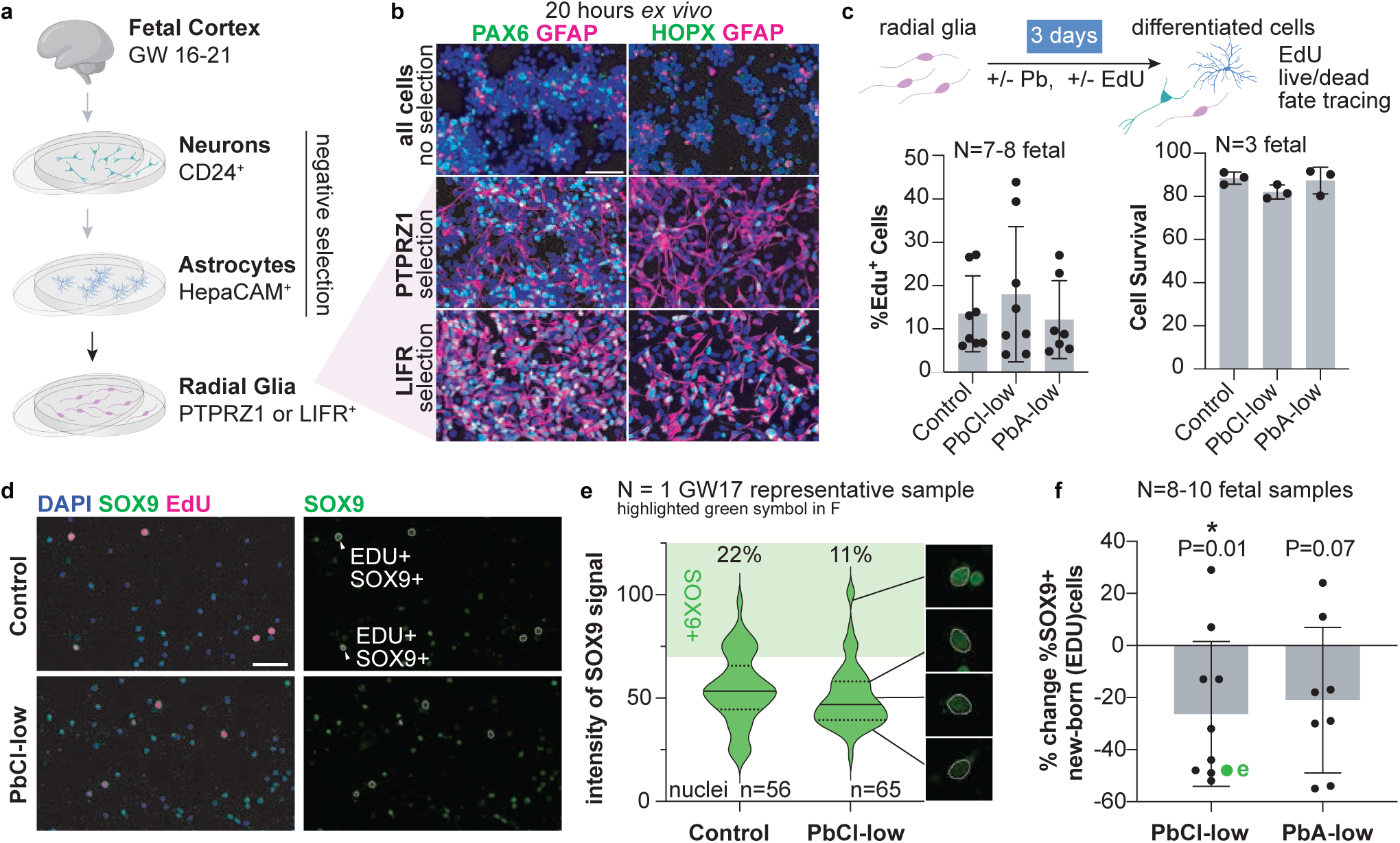
Fetal radial glia exposed to Pb exhibit inhibited astrocyte differentiation and increased H3K27me3 repressive mark. **a**, Radial glia were immunopanned (LIFR or PTPRZ1) from dissociated fetal cortices following depletion for CD24+ neurons and HepaCAM+ astrocytes. **b**, Representative images from dissociated cells with no immunopanning selection or immunopanned PTPRZ1 or LIFR cells. Cells were immunostained with PAX6 or HOPX (green) and GFAP (magenta) to quantify radial glia enrichment. Scale bar is 50 µm. **c**, 2D cultures of fetal radial glia were exposed to low doses on PbCl_2_-low or PbA-low for 3 days. EdU was added to quantify proliferation in culture (N=7-8 fetal samples) and live-dead imaging was used to quantify survival (N=3 fetal samples). Each point represents data from an individual fetal tissue sample. **d**, New-born cells were identified by EdU (magenta) and stained for SOX9 (green). Images shown are from a representative GW17 fetal sample. Scale bar is 50 µm. **e**, SOX9 immunoreactivity quantified for individual nuclei. The SOX9+ threshold is shaded in light green. Examples of nuclei across a spectrum of SOX9 intensity are shown in the right inset panels. **f**, Percent change in the %SOX9+ newborn cells relative to control for N=8-10 individual fetal samples. Data corresponding to the representative sample in **e** is highlighted in green.

Because we observed significant changes in SOX9+ / EdU+ progeny in the presence of Pb, we wondered if there were other direct upstream modulators of SOX9-dependent astrocyte fate acquisition. NOTCH signaling is a known regulator of *SOX9* expression and our previous GSEA analysis from acutely Pb-exposed whole organoid RNA-Seq suggested decreased NOTCH pathway activity (Extended Data Fig. 1f). We therefore measured protein levels of both full length and NOTCH activated NOTCH intracellular domain (NICD) in PTPRZ1 cell protein lysate by immunoblot (Extended Data Fig. 4d). Using the ratio of NICD/FL protein as a readout of NOTCH activity, we found decreased NOTCH signaling activity in Pb-high (500 µM) conditions, but no consistent change with Pb-low (500 nM). Importantly, NOTCH pathway activity is repressed by PRC2^46^, which is consistent with our previous observations of augmented PRC2 activity in the presence of Pb.

In our acute exposures to Pb-high (500 µM) we observed increases in oxidative phosphorylation genes, while chronic, Pb-low (500 nM) was associated with an increase in glycolysis genes. To examine whether Pb-induced changes in cellular metabolism could contribute to the push toward neurogenesis we isolated neurons, astrocytes and radial glia from primary fetal cortex (GW18-21) and performed Seahorse extracellular flux oximetry assays (Extended Data Fig 4f-h). Non-mito oxygen consumption was significantly decreased in the neurons (P=0.03, one-way t-test) and neurons were significantly different than astrocytes (P=0.0165 by Brown-Forsythe ANOVA test, P=0.039 Dunnett’s T3 multiple comparisons test). Although the data suggest that the three cell types had different bioenergetic responses to Pb, we were unable to detect consistent changes in radial glia bioenergetics that could have a causal role in cell fate bias (Extended Data Fig 4f-h).

### PbCl_2_ exposure increases intermediate progenitors and decreases glial progenitor cells

Pb exposure shifted how radial glia differentiate in culture, ultimately reducing the number of cells expressing the astrocyte commitment marker SOX9 (Fig. 4f). While this ICC-based approach enabled us to restrict our analysis to newly generated cells using EdU labeling, it limited our ability to assess cell identity to only a small number of markers. To more precisely define the identity of newborn cells, we used a viral lineage tracing approach called CellTag^47^, which enabled us to simultaneously track both clonal cell lineage commitment and single-cell transcriptomic signatures. Celltag is a lentiviral-based transcribed barcode approach that includes ∼10^12^ unique barcodes within the 3’UTR of a GFP sequence to allow identification of clonal cells^47^. We infected primary PTPRZ1-enriched fetal cells (GW16-20) with the lineage tracing virus (MOI 3) for 24 hours and then exposed the cells to control or Pb conditions for 6-7 days. After exposure, we isolated GFP+ cells by FACS and captured for 3’ 10X single-cell sequencing (Fig. 5a). We prepared both 10X single cell libraries and side libraries that amplify the GFP UTR containing the CellTag barcodes.

**Fig. 5.**
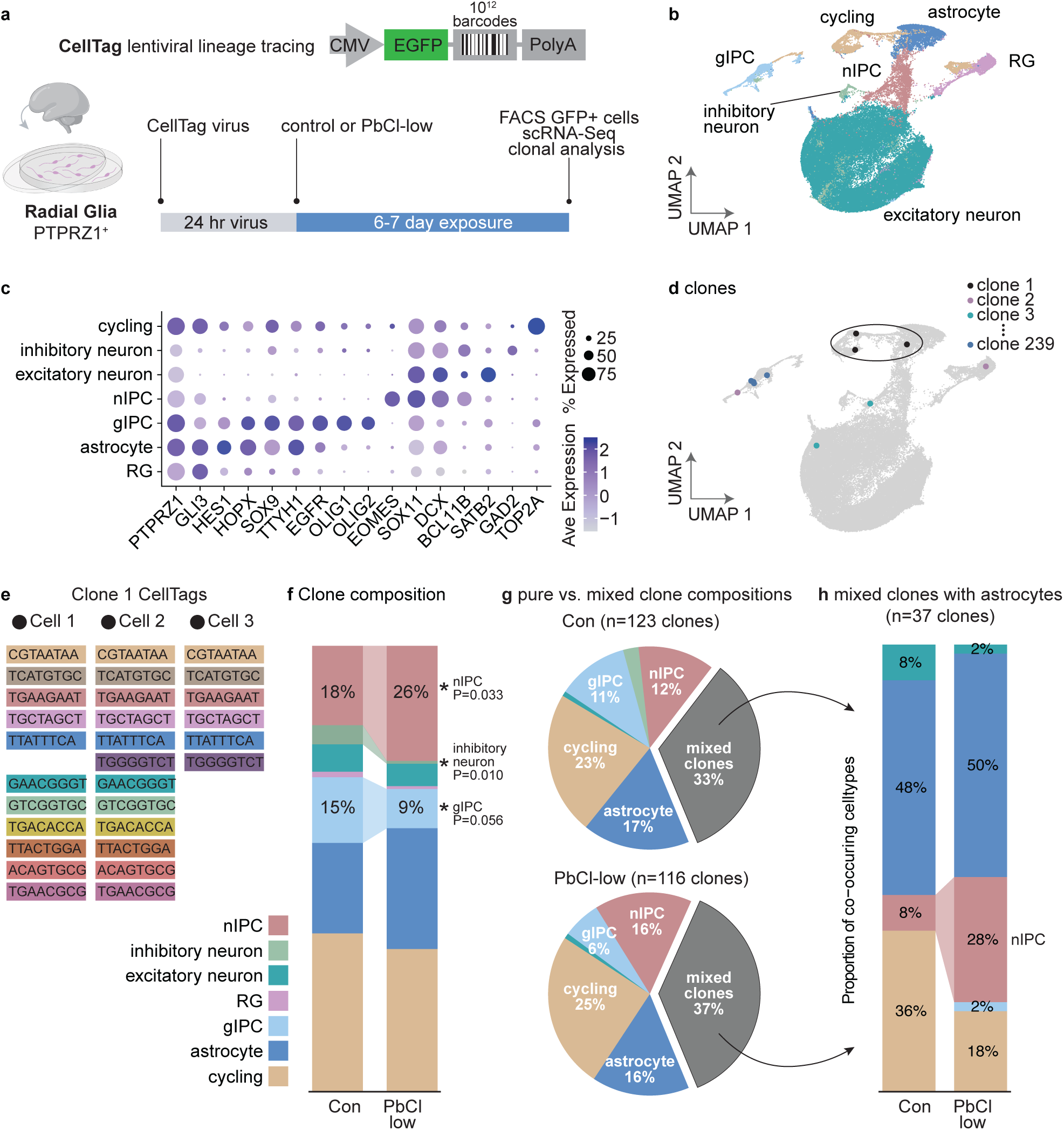
Clonal lineage tracing of PTPEZ1+ radial glia exposed to Pd demonstrate a bias towards neuronal intermediate progenitors and fewer glial progenitors. **a**, PTPRZ1 cells were immunopanned from human primary cortex GW16-20, exposed to CellTag lentivirus for 24 hours, then PbCl_2_-low (500nM) for 6-7 days. GFP+ (CellTag) cells were FACS isolated and captured for single-cell and clonal analysis. **b**, UMAP showing PTPZR1-derived cell clusters after 8 days in culture. **c**, Selected gene markers for cel clusters. **d**, Example CellTag clones are show overlayed on the UMAP. **e**, Clone 1, which contains 1 astrocyte cell and 2 cycling cells, has 5 barcodes with at least 2 UMIs shared across all three cells. **f**, Clone composition averaged for control and PbCl_2_-low exposed cells. P-values are from a one-way t-test of the percent change of PbCl_2_-low for N=3 fetal samples. **g**, Clones were composed of either “pure” single-cell types or mixed populations of cells. **h**, The composition of astrocyte-containing mixed clones normalized by the total cells in the clone group.

From N=3 fetal samples (GW 16, 18, 20), a total of 86,308 cells passed QC and we identified all major cortical cell classes including excitatory neurons, intermediate progenitors, inhibitory neurons, glial progenitor cells, and radial glia (Fig 5b-c, Extended Data Fig 5a-b). We observed multiple classes of progenitors including radial glia (RG), neuronally committed intermediate progenitor cells (nIPCs), and glial intermediate progenitor cells (gIPCs). Pb exposure increased the proportion of radial glia in the data set by 33% (N=3 hiPSC lines, one-sample t-test P=0.0146) whereas all other classes of cells were either unchanged or decreased (Extended Data Fig 5a-b). To focus our analysis to actively expanding cells, we performed clonal analysis using CellTag barcodes. We identified 239 clones using strict cutoffs (minimum of 2 UMIs per CellTag, minimum of 2 unique CellTags per cell, jaccard threshold of 0.6 (Extended Data Fig 5c) to ensure robust clonal relationships between cells. Several example clones are highlighted in Fig 5d, along with the sequenced CellTags for each cell within clone 1 (Fig. 5e).

Comparing the clone composition of control vs Pb-exposed cells revealed several differences: an increase in nIPCs, from 17.8% in control to 25.6% in Pb-exposed clones (P=0.033), a decrease in inhibitory neurons from 4.3% to 0.6% (P=0.010), and a trending decrease in gIPCs from 14.7% to 8.75% (P=0.056) (Fig 5f). Although nIPCs increased within the clonal populations, which are restricted to actively expanding cells, this was not the case in the dataset overall, where nIPCs were 4.6% in control and 4.5% in PbCl-low. This is possibly a reflection in decreased neuronal survival in a population not selected for actively dividing cells (Extended Data Fig 5b). Interestingly, although radial glia numbers were increased upon Pb-exposure in our single cell data set overall, this population was largely absent from our clones, suggesting that our captures were biased towards terminal differentiations or possibly impacted by CellTag infection bias.

We next examined how Pb exposure affected clones composed of ‘pure’ single cell types vs mixed populations (Fig. 5g). Pb-exposed cultures had fewer pure gIPC clones compared to control (11% of PbCl_2_ clones vs 6% of control clones), and more pure nIPC clones compared to controls (16% PbCl_2_ clones vs 12% control clones) (Fig. 5g). We reasoned that clones with multiple cell lineages may be reflect a shift in differentiation, so we performed a similar analysis focused on mixed lineage clones. There were n=37 mixed clones containing astrocytes and these clones contained 3.5 times more nIPCs (8% control vs 28% PbCl_2_) (Fig. 5h), possibly reflecting a shift toward the neuronal lineage over the course of the 6-7 day Pb exposure.

### Transient PbCl_2_ exposure signature persists for 7 weeks in mouse transplantation model

We next asked whether Pb-induced changes in gene expression and cell fate in developing neural progenitors persist even after a transient exposure period. This is difficult to assess in 2D monoculture because while fetal cells grow robustly in culture over short time durations (days to weeks), they begin to lose integrity over longer timescales of weeks to months. One approach to sustain long-term cultures of fetal cells is xenotransplantation, in which cells are transplanted and engrafted into an *in vivo* host tissue such as the mouse brain. We used this method to test whether transient Pb exposures elicit long term effects that persist in fetal cells after transplantation into non-exposed immunodeficient mice. Prior to transplantation, we infected primary fetal PTPRZ1 immunopanned cells (N=1, GW 18) with CellTag lentivirus. We simultaneously exposed these cells to either PbCl_2_-low or NaCl for 10 days and transplanted them into subcortical regions of P2 mouse brains (N=2 NaCl, N=3 PbCl_2_-low into 5 pups from a single litter) where they engrafted and migrated throughout the parenchyma (Fig 6a-b). After 7 weeks we dissected brain tissue, dissociated the tissue into single cells, and cleaned the cells with MACS myelin removal beads. We pursued several steps to remove mouse cells from the cell suspension prior to single cell capture, including MACS mouse cell depletion beads and FACS for GFP+ cells (human). Following these steps, we captured GFP+ cells using our standard 10X capture protocols.

**Figure 6.**
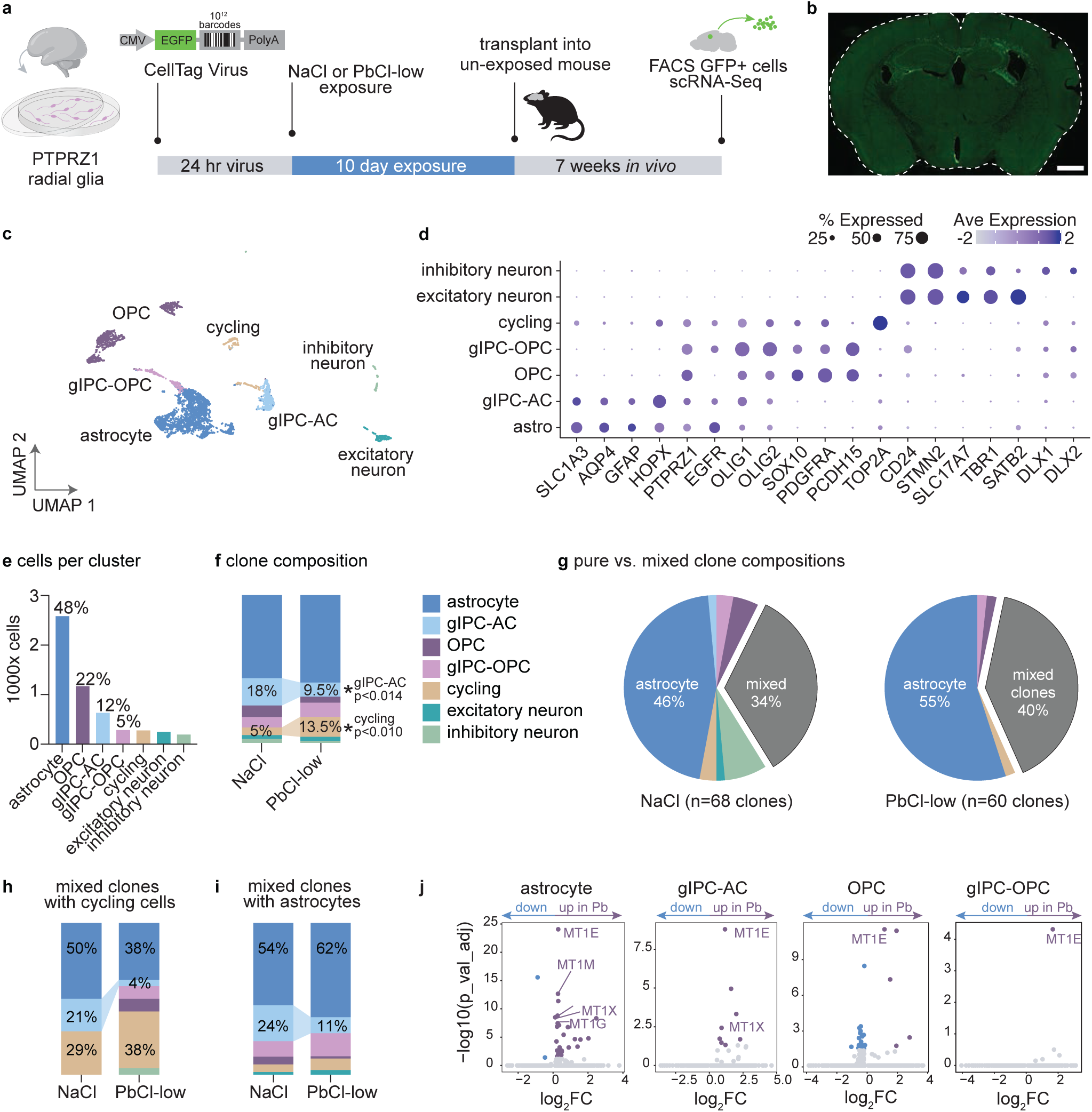
Fetal PTPRZ1+ radial glia show persistent Pb transcriptional signature 7 weeks after exposure. **a,** PTPRZ1 cells were immunopanned from a GW18 fetal brain, infected with CellTag VI virus, exposed to PbCI2-low or NaCl for 10 days, then transplanted into 5 unexposed P2 pups. After 7 weeks, fetal cells were isolated from the brain for single cell sequencing, **b,** By 6 weeks, the cells predominantly migrated into white matter tracts. Scale bar is 1000 µm. **c,** Human PTPRZ1-lineage cells isolated from the mouse brain and sequenced using 10X visualized on a UMAP. **d,** Selected markers demonstrating cell­specificity of each cluster, e, The number of cells (y axis) per cluster (x axis) and the proportional percentage of each cluster is shown above the bars, **f,** Clone composition for each treatment group, **g,** Clones were composed of either “pure” single-cell types or mixed populations of cells, **h,** The composition of mixed clones containing cycling cells normalized by the total cells in the clone group, **i,** The composition of mixed clones containing astrocyte cells normalized by the total cells in the clone group, **j,** Cluster-specific transcriptional response to Pb.

To ensure that we specifically analyzed Pb effects on human cells and not residual contaminating mouse cells, we aligned the single cell data to both human (GRCh38) and mouse (GRCm39) genomes using Cell Ranger. We used the number of UMIs assigned to either mouse or human genes in each cell to assign to a species (Extended Data Fig 6a). Cells exceeding UMI counts in the 10^th^ percentile for both species were assigned as multiplets and not included in subsequent steps. Only human cells with less than 10% human mitochondrial genes and a minimum of 800 genes (nFeature) were included in downstream analysis.

The fetal isolated PTPRZ1-lineage cells were transplanted into a postnatal brain, which is a highly pro-gliogenic environment. In line with this, we observed that most of the transplanted cells differentiated into macroglia: 48% astrocytes, 22% oligodendrocyte precursor cells (OPCs), and 12% astrocyte progenitor cells (GIPC-AC progenitors). The remaining cells had gene expression profiles consistent with 5% gIPC-OPC progenitor, 5% cycling cells, 4.6% excitatory neurons, and 3.6% inhibitory neurons (Fig 6c-e). Within CellTag clones, GIPC-AC cells were reduced from 18.2% to 9.5% (P=0.014, t-test) and cycling cell populations increased from 5.2% to 13.5% (P=0.010, t-test) in the Pb-exposed group (Fig 6f, Extended Data 6b), again reflecting the lineage biases we observed in previous exposure paradigms. We observed an increase in both pure astrocyte clones and mixed clones in the Pb-exposed clones (Fig. 6g). When we focused on mixed clones containing cycling cells, we observed a marked decrease in both GIPC-AC and astrocytes (Fig. 6h). In astrocyte containing clones, there was again a decrease in GIPC-AC cells (Fig. 6i).

Remarkably, even 7 weeks after Pb exposure, astrocytes, OPCs, and progenitor populations in the mouse brain retained elevated expression of metallothionein genes, especially MT1E. This upregulation was specific to PbCl₂-exposed cells, and we did not observe the same changes in neurons (Fig 6g, Extended Data Fig. 6c). At the transcriptomic level astrocytes derived from Pb-exposed progenitors exhibited an increase in metallothioneins, mitochondrial genes (MT-CO3, MT-CO1), neural adhesion proteins (CNTN1, CNTNAP2, OPCML), and cytoskeleton components (GFAP, TUBB3, TMSB10). Interestingly, though the metallothionine signature persisted, we did not observe transcriptional changes reflecting oxidative stress or antioxidant response, except for a modest increase in thioredoxin (TXN) in astrocytes.

## Discussion

We established an exposure strategy for cortical organoids and found that Pb exposure decreases the number of astrocytes, while increasing excitatory neurons. In primary radial glia cultures, Pb acts directly on neural progenitors to alter lineage commitment and single-cell lineage tracing revealed an increase in nIPCS at the clonal level following Pb exposure. Our data showed an increase in neurogenesis and this finding is consistent with two previous organoid exposure studies wherein Pb exposure (1) increased neuronally committed intermediate progenitors in hESC-derived organoids^22^ and (2) exhibited higher expression of the neuronal marker TUJ1 at the expense of mature neurons in hiPSC-derived organoids^23^.

Together, these data suggest that toxicologically relevant Pb exposures can bias lineage commitment towards neurogenesis at the expense of astrogenesis and radial glia may be particularly vulnerable to Pb exposure during their transition from neuronal to glial fates, often referred to as the ‘gliogenic switch.’

In both our organoid single-cell and time course studies we observed an increase in gene signatures associated with the epigenetic regulator PRC2 following Pb exposure. This protein complex deposits the repressive H3K27me3 to regulate gene expression programs. In Pb-exposed radial glia, we measured an increase in H3K27me3, directly implicating an increase in PRC2 activity or a decrease in demethylation. The core PRC2 complex associates with several accessory proteins including MTF2 to form PRC2.1 or JARID2 to form PRC2.2. It is thought that the accessory proteins act as rheostats for cell fate decisions, modulating the activities of the core PRC2 complex to be appropriate for different cell lineages^48^. MTF2 is particularly interesting as it is known to regulate canonical WNT signaling^49^ and is also highly expressed in early cortical neurons and follows the expression patterns of SOX4 and SOX11^50^. Additionally, the full name for MTF2 is ‘Metal Response Element Binding Transcription Factor 2,’ because it contains high sequence similarity with the ‘metal response element’ of the canonical metallothionine MT1A. Whether MTF2 retains metal response functionality and links Pb exposure to PRC2 activity is an intriguing possibility.

PRC2 has been previously implicated in cell fate, particularly in neurogenesis and gliogenesis, across rodent^44^, bird^46^, and human^50^ systems. In the murine cortex, loss of PRC2 narrows the neurogenic window leading to precocious astrogenesis^44^. In chicks, loss of PRC2 in oligodendrocyte lineage cells caused aberrant expression of astrocyte genes and suppressed oligodendrocyte differentiation, but this was rescued by inhibition of Notch signaling^46^. In hiPSC organoids, PRC2 has also been shown to be an epigenetic barrier to neuronal maturation, where the release of PRC2 and H3K27me3 was associated with accelerated neuron maturation and increased astrogenesis^50^. These three previous studies demonstrated that loss of PRC2 increases astrogenesis, whereas our study demonstrated that an *increase* in H3K27me3 correlated with an increase in neurogenesis and concomitant decrease in astrogenesis. Our data, in the context of these studies, provides further evidence for an important role for PRC2 in cell fate decisions.

Previous work suggests that cell stress and hypoxia both promote neuronal cell fate and these factors may also contribute to the Pb exposure associated promotion of neurogenesis and suppression of astrogenesis. There was a hypoxia gene signature in astrocytes, neurons, outer radial glia and ventral radial glia after 3-week exposure to PbA-low. It has been previously reported that metals, including Pb(II), can induce “pseudohypoxia” where HIF-1α accumulates under normoxic conditions^51^. In addition to a pseudohypoxia signature, there was a strong antioxidant response both by gene expression and quantification of glutathione in organoids. The neuroprotective ratio of reduced glutathione was sufficient to protect the cells from loss of redox potential even at Pb-high (500 µM), suggesting that the organoids are resilient against oxidative stress. It is unclear whether this is an artifact of organoid culture or whether intact primary tissue would also exhibit high cellular survival with high Pb exposures. In our hands, fetal sections (300 µM thick) are also resilient to high lead exposures compared to more vulnerable 2D cultures. The real-world relevant exposures of Pb-low (500 nM) showed greater than 70% cell survival in both organoids and fetal cells compared to ∼90% survival in unexposed controls.

To model transient, human Pb exposure, we treated fetal radial glia with PbCl₂ or NaCl for 10 days before transplanting them into the postnatal mouse brain. Remarkably, even after 7 weeks *in vivo*, Pb-exposed cells retained a strong metallothionein gene signature—one of the most robust transcriptional changes we observed. Using CellTag lineage tracing to isolate clonally expanding cells, we found a decrease in GIPC-AC cells and increase in cycling cells, which is remarkable considering these cells differentiated under the pro-gliogenic pressures of a postnatal mouse brain. For this experiment, we isolated transplanted cells from the white matter, where they were most abundant, to maximize retrieval yield and reduce the time from dissociation to cell capture. These steps may have biased our findings. It is possible that cells that migrated to other regions exhibit different cellular composition or expression profiles. Additionally, this experiment was limited to a single GW18 fetal sample due to the challenge of compatible timing of human tissue procurement and mouse litters births. However, even with the limited sample size, this experiment demonstrated a persistent transcriptional signature in astrocytes and OPCs even 7 weeks after exposure.

To examine Pb neurotoxicity in the context of the human genome, we leveraged hiPSC cortical organoids and complemented these findings with primary fetal cortex tissues. Primary human tissues are challenging due to sample-to-sample variability (e.g., genetics, age, sample integrity) and limited availability, whereas the organoid approach offers limitless expansion of hiPSC lines that enable rigorous replication and region-specific differentiation. There remain limitations to organoid model systems that are relevant to this study: cortical organoids typically contain few inhibitory neurons and OPCs, limiting our ability to assess the effects of Pb on these populations. Future studies could address this by incorporating subpallial regions to examine whether Pb exposure alters the generation of inhibitory neurons or OPCs. While this limitation is inherent to region-specific organoids, it also enables focused investigation of a critical cell fate decision: the divergence of neural intermediate progenitors (nIPCs) towards excitatory neurons versus astrocyte precursor cells (APCs) towards the astrocyte lineage.

Prenatal Pb exposure is associated with cognitive and developmental deficits, but the cellular and systems level outcomes underlying these observations remain unknown. Our data suggest that one potential consequence of prenatal Pb exposure is a prolonged neurogenic period and delayed astrogenesis onset, associated with increased H3K27me3, which is deposited by chromatin modifier PRC2. Cell fate bias was demonstrated in both primary *ex vivo* radial glia cultures and human induced pluripotent stem cell-derived 3D organoids, suggesting translatability to real-world exposures. Understanding how Pb disrupts excitatory neurogenesis is particularly relevant to modeling neurodevelopmental disorders characterized by excess excitatory neuron production. For example, organoids derived from donors with idiopathic autism spectrum disorder have shown increased neurogenesis^52,53^, including elevated DLX2 expression and other genes overlapping with our observed Pb-associated transcriptome. Future studies could combine the exposure paradigms established in the current study with patient-derived hiPSC derived organoids to model brain development within the context of real-world disease gene-environment interactions^27^.

Our findings highlight a previously underappreciated sensitivity of radial glia to Pb toxicity, even in the context of transient, low level exposures, which induced persistent transcriptional changes that could potentially alter cell maturation and function for months. Our data suggest that Pb shifts lineage trajectories—promoting neurogenesis at the expense of astrogenesis—offering a compelling mechanistic link between epigenetic modulators and the risk for cognitive delay and neurodevelopmental disorders. This insight into the molecular mechanisms underlying toxicant associated impairments will guide future efforts to develop actionable prevention strategies and deepen our understanding of the etiologies underlying cognitive and behavioral impairments associated with Pb exposures.

## Methods

### Cortical Organoids

Cortical organoids were formed as previously described^25,33^ from human induced pluripotent stem cells (hiPSCs). All hiPSC lines were tested for mycoplasma and genomic integrity (karyotyping). To begin, 80-90% confluent iPSC colonies were detached from the plates with Accutase (VWR, 10761-312) and were formed into 3D spheroids in Aggrewell 800 Plates (24-well) (STEMCELL, 34811) overnight. Spheroids were grown in neural induction media for 6 days then switched to Neurobasal-A (ThermoFisher, 10888022) supplemented with B-27 (ThermoFisher, 12587010) and Penicillin/Streptomycin (VWR, 16777-164). Patterning was achieved with Dorsomorphin (Sigma, P5499-25MG, 5 µM) and SB-431542 (Selleck Chemicals, S1067, 10 µM) on days 1-6, EGF (R&D Systems, 236-EG) and FGF (R&D Systems, 233-FB) for days 7-15, then BDNF (PeproTech, 450-02-1mg) and NT-3 (R&D Systems, 267-N3-005/CF) for days 16-42. Thereafter, organoids were fed with neurobasal A with B-27 and Penicillin/Streptomycin. Human induced pluripotent stem cells 8858.3, 2242.1 and 1363.1 were previously generated from Sendai viral reprogramming of human fibroblasts acquired from healthy control patients at Stanford University. Line C4.1 was obtained from Coyne Scientific and 0524-1 was sourced from the Pasca Lab (Stanford University). Additional hiPSC line metadata are listed in the “Source_Data” file.

### Human Tissues

All human tissue samples were obtained in compliance with policies outlined by the Emory School of Medicine IRB office. All experiments were conducted in compliance with all applicable federal, state, or local laws including regulatory and informed consent processes. HFT from consenting individuals was procured via overnight shipping through a third-party. Cortical regions were identified by the gross morphology stereotypical of mid-gestation cortical tissue (curved exterior, faint appearance of gyri and sulci, density along the apicobasal axis of a coronal section). All tissue samples are listed in the “Source_Data” file.

### Tissue/Organoid Dissociation and Immunopanning

Fetal tissue was dissociated into a single cell suspension using previously published methods^54^ and this was adapted for organoids by decreasing volumes and omission of filtering steps. Briefly, tissue was chopped using a scalpel and incubated in papain (7.5 units/mL for HFT and 25 units/mL for organoids) at 34 °C for 45 min before rinsing with a protease inhibitor solution (ovomucoid) and mechanical trituration. Dissociated cells were immunopanned to enrich specific cell types from organoid or primary fetal brain tissue as previously described^19,25^. Astrocytes were isolated using mouse anti-HepaCAM (R&D systems, MAB4108), neurons were isolated using mouse anti CD24 (Miltenyi Biotec, 130-108-037) or mouse anti-Thy1 (BD Biosciences 550402), and radial glia were isolated using mouse anti-PTPRZ1 (Santa Cruz Biotechnology, sc-33664). The secondary antibody was Goat anti-mouse IgG+IgM (H+L) (Jackson ImmunoResearch, 115-005-044). All immunopanned cells were plated on poly-D-lysine (Sigma-Aldrich P6407) coated plastic coverslips and fed with a 50:50 Neurobasal/DMEM based serum-free medium^19^ supplemented with Penicillin/Streptomycin, 1 mM Sodium Pyruvate, 2mM L-glutamine, 5µg/ml NAC, 1 µg/mL transferrin (Sigma T-1147), 1 µg/mL BSA, 0.16 µg/mL putrescine (Sigma, P5780), 2 nM progesterone (Sigma P8783), 0.4 ng/mL sodium selenite (Sigma, S5261), and 5 ng/mL HBEGF.

### Pb Exposures

Organoids were fed with media supplemented with Pb acetate trihydrate (Sigma-Aldrich 316512-5G) or PbCl_2_ (Sigma-Aldrich, 203572). Control organoids were fed with media alone or media supplemented with sodium acetate (Sigma-Aldrich, 229873), or sodium chloride (Sigma-Aldrich, S3014). For proliferation experiments, organoids or cells were fed with media with supplemented with 10 µM EdU (Molecular Probes, A10044).

### Leadmium Live Imaging

Lead uptake was visualized with leadmium fluorescent sensor (Thermo Fisher, A10024) according to the manufacturer’s instructions. Fetal or organoid cells were immunopanned (astrocytes or neurons), plated on PDL-treated coverslips, and fed with neurobasal media. Cells were incubated with saline with leadmium for 30 minutes. Baseline was recorded for 20 min to calculate DF/F. Live imaging was performed on a Keyence fluorescence microscope (488 nm excitation) with images taken every 2 min.

### Pb quantification

Single organoids were collected into dry tubes, massed and frozen. Frozen tissue was heat digested with concentrated nitric acid before the addition of dilute nitric acid and a solution containing the internal standard (ISTD) iridium. The samples were then analyzed on a 7700 Series ICP-MS (Agilent Technologies, Santa Clara, CA) alongside a six-point calibration curve, removing polyatomic spectral interferences with a dynamic collision reaction cell. Concentrations of Pb were derived from an equation defining the relative response of the element to the ISTD response across the calibrant concentration range. Quality control materials and blank samples were analyzed concurrently with organoid samples to ensure quality measurements were obtained.

### GSH/GSSG quantification

Reduced and oxidized glutathione were measured in organoid tissues by Creative Proteomics. Briefly, for each analytical sample 3 organoids were pooled (totaling 3-9 mg/pooled sample). The pooled tissue was homogenized in water and methanol, then spiked with an internal standard solution of reduced glutathione (glycine-13C2,15N). Then 10 μL of each standard solution or each resultant sample solution was injected into a HILIC column (2.1 × 100 mm, 1.7 μm) for UPLC-MRM/MS runs with (+) ion detection on an Agilent 1290 UHPLC system coupled to an Agilent 6495C QQQ MS instrument, with the use of 0.1% formic acid in water and 0.1% formic acid in acetonitrile as the mobile phase for gradient elution (85% to 25% B over 10 min) at 0.3 mL/min and 40 °C.

### Bulk RNA-Seq

RNA was extracted using RNeasy Micro kit (Qiagen 74004) according to the manufacturer’s protocol. For whole organoids, library preparation was performed with NEBNext Ultra II RNA (non-directional) with PolyA selection. For immunopanned cells, libraries were prepared using the SMART-Seq® HT PLUS Kit (Takara Bio USA, R400748 & R400749). Sequencing was performed using Illumina 2×150 platforms to a depth of 40M PE reads (20M each direction). Fastq files were mapped to hg38 using STAR aligner and FeatureCounts to generate raw count matrices. The counts were normalized using edgeR TMM and differential gene expression was performed using Limma.

### 10X Single-Cell Sequencing

All dissociation steps were performed in the presence of Protector RNAse inhibitor (Sigma 03335402001), transcription inhibitor actinomycin D (Sigma-Aldrich; A1410), and translation inhibitor anisomycin (Sigma-Aldrich, A9789) based on recommendations in PMID: 35260865. Cells were incubated with CellPlex CMOs on ice for 5 min, washed, then incubated with 50 nM calcein-AM and 4 uM ethidium homidimer-1 from the LIVE/DEAD™ Viability/Cytotoxicity Kit (Thermofisher Scientific, L3224) prior to cell sorting. Live (calcein+, ethidium-) cells sorted on a BD Facs Aria II. Cells were sorted into 4% BSA with RNAse inhibitor. For cell CellTag lineage tracing, GFP+ cells were isolated by FACS before capture and library preparation. Single-cell captures were performed using the 10X chromium platform. For D150 organoids, dissociated cells labeled with CellPlex CMOs (10X 1000261), FACS purified, counted and loaded on the chromium chip to achieve target recovery of 10,000 cells.

Capture and library prep were completed according to 10X protocols for Chromium Next-GEM single cell 3’ reagent kit v3.1 (dual-index) (PN-100268).then were captured on a Chromium Next GEM Chip G (10X 1000120). Libraries were prepared using Chromium Next GEM Single Cell 3′ Kit v3.1 (10X 1000268) and Dual Index Kit TT Set A (for Gene Expression Libraries) or Dual Index Kit Set A NN (for sample multiplexing) (1000243). Sequencing was performed by Admera Health on a NovaSeq S4 2×150 to achieve 50-100k reads per cell. CellPlex library was sequenced at a depth of 5-10k reads per cell.

Cells were demultiplexed using Cell Ranger. Analysis was performed in R Studio using Seurat. For quality control, cells with <8% mitochondrial genes were excluded. Cells were integrated and normalized using SCT transform to generate a UMAP and clustering based on based on shared nearest neighbors. All featureplots, dot plots, and differential expression are based on non-transformed RNA counts. Differential expression was performed using pseudobulk data aggregation and Findmarkers DeSEQ2 or MAST. Heatmaps are based in scaled RNA counts data.

### CellTag Lineage Tracing

Immunopanned-PTPRZ+ fetal brain cells were treated with V1 or V3 CellTag virus (Addgene, 115643-LVC or 115645-LVC) at MOI 3 for 24 hours. Infected cells were purified by FACs before single-cell capture. For demultiplexing, side-libraries were generated using primer sequences modified from Jindal 2024^55^. The forward primer adds a P7 sequence, 8bp index, and TruSeq Read2 sequence, it relies on eGFP specific portion to anneal to 3’ end of eGFP going forward into the CellTag barcode and the 3’UTR. The index is any 8bp sequence that doesn’t interfere with other multiplexing indices. The reverse primer adds a P5 sequence and uses a partial TruSeq Read1 sequence to anneal to the Read1 adaptor added to your cDNA during “Step2.2 cDNA amplification” in the 10x library prep protocol.

### Fetal Xenotransplantation

Immunopanned-PTPRZ+ fetal brain cells (GW18) were plated in 6-well plates and allowed to recover. On 2 DIV, cells were infected with V1 CellTag virus (Addgene 115643-LVC) or CMV-GFP virus. On 3 DIV, media was supplemented with either 1000 nM NaCl or 500 nM PbCl_2_-low, which were added at each media change for 10 days. On 13 DIV, 200,000-400,000 PTPRZ1+ cells into postnatal day 2 Rag2 knockout pups (JAX, strain #008449)^56^. After 40 days, CMV>GFP mice were sacrificed for IHC, and we visualized cells were preferentially migrated into the white matter. After 47 days, CellTag-infected xenograft mice were sacrificed. The corpus callosum, along with small proportions of deep-layer cortex, dorsal striatum, dorsal thalamus, and hippocampus, was dissected, and the tissue was dissociated into a single cell suspension using papain digestion following a previously published protocol^57^. Cells were then purified using MACS Myelin Removal Beads II (Miltenyi Biotec, 130-096-433) and MACS Mouse Cell Depletion Beads (Miltenyi Biotec, 130-104-694) before isolated GFP+ cells via FACS. Purified cells were captured via GEM-X (PN-1000691) with 4-7,000 cells loaded per well.

### RNA-Seq analysis

Gene Set Enrichment Analysis (GSEA) was performed using the R package clusterProfiler^58^ and the Molecular Signatures Database (gsea-msigdb.org) with either Hallmark (H) or curated (C2) gene sets. Gene Ontology was performed with the R package gProfiler2^59^. Weighted Correlation Network Analysis^60^ (WGCNA) was performed in R to identify modules of genes that correlate over the course of Pb exposure. Enrichr^61^ was used for ChIP enrichment analysis^62^ (ChEA).

### Organoid IHC and fetal ICC

Organoids and 2D cells were washed with PBS and fixed using 4% Paraformaldehyde Aqueous Solution (EMS, 15710). Organoids were embedded in Optimum Cutting Temperature (OCT) Compound (VWR, 25608-930) and cryosectioned to 12-20 μm and added to glass slides. For cell death and proliferation, we performed TUNEL (Invitrogen, C10618) or EdU (Fisher Scientific, C10640) staining with DAPI (Vector, H-1500-10). Slides were then imaged on Keyence fluorescence microscope at 40X magnification. Cells were counted using Keyence analyzing software.

Fetal cells grown on coverslips were stained with 1:300 anti-SOX9 (R&D Systems, AF3075, 48 hours), 1:1000 anti-TUJ1 (Novus, NB100-1612 or BioLegend, 801201), anti-GFAP(BioLegend, PCK-591P or Agilent Dako, Z0334), anti-PAX6 (Cell Signaling Technology, 60433S), anti-HOPX (Sigma, HPA030180), or 1:800 H3K27me3 (Cell Signaling Technology, 9733S). We used Alexa Fluor 647 Click-iT™ EdU Cell Proliferation Kit (Invitrogen, C10340) to identify newborn cells.

### Live/Dead Quantification

Cells were incubated with 50 nM calcein-AM and 4 uM ethidium homidimer-1 from the LIVE/DEAD™ Viability/Cytotoxicity Kit (Thermofisher Scientific, L3224) and coverslips were transferred onto slides. Slides were imaged on Keyence fluorescence microscope at 40X magnification. Cells were counted using Keyence analyzing software.

### Organoid Proliferation Quantification with EdU/Flow

Organoids were fed with media with supplemented with 10 µM EdU +/-PbCl_2_ for 3 weeks (D130-D150). On D150, organoids were dissociated into a single suspension as described above. Single cells were then fixed and stained for Edu using Click-iT™ EdU Alexa Fluor™ 647 Flow Cytometry Assay Kit.

### Immunoblots

Protein lysates were prepared by incubating the organoids in protein extraction buffer (150 mM NaCl, 20 mM Tris–HCl (pH 7.5), 5 mM MgCl2, 1% NP-40, supplemented with protease and phosphatase inhibitor cocktail (Sigma-Aldrich, PPC1010; dilution 1:100)) on ice for 10 min, followed by probe tip sonication (Misonix Sonicator 3000, 20–30 pulses). Protein concentrations were determined using a BCA protein assay (Thermo Fisher Scientific, 23227). Equal concentrations of protein lysates (20 µg) were loaded on 4–12% Bolt Bis–Tris Plus Gels (Invitrogen, NW04120BOX) separated by SDS-PAGE (Invitrogen, NW04120BOX) and electro-transferred onto nitrocellulose membranes (Invitrogen, IB23001) using the iBlot 2 Dry Blotting system. Membranes were blocked in 5% BSA in TBST (150 mM NaCl, 20 mM Tris (pH 7.6), 0.1% Tween 20) and probed with the appropriate primary antibody overnight at 4 °C. The following antibodies were used: actin (Sigma-Aldrich, MAB1501; dilution 1:2,000), Notch/NICD (R&D Systems AF3647). Secondary anti-mouse and anti-rabbit antibodies conjugated to horseradish peroxidase (Invitrogen, 31430 and 31460, respectively) were used at dilutions of 1∶10,000 and incubated with the membranes at room temperature for 60 min. Proteins were visualized by enhanced chemiluminescence (Thermo Fisher Scientific, 34577) and imaged using a Bio-Rad ChemiDoc Imaging System. Protein signals were quantified using Image Studio Lite 5.2.5 Quantification Software (LI-COR Biosciences) or FIJI.

For H3K27me3 and H3 quantification, histones were acid-extracted from whole organoids. Briefly, organoids were mechanically lysed by resuspension in Triton extraction buffer (TEB) containing 0.5% Triton X-100 and supplemented with protease inhibitors (7.5 μM aprotinin, 0.5 mM leupeptin, 250 μM bestatin, and 25 mM AEBSF–HCl). Cell lysates were centrifuged at 6,500 × *g* for 10 minutes at 4°C. Histones were then acid-extracted from the resulting pellet using TEB supplemented with 0.8 M HCl at a 1:1 ratio. After 20 minutes of incubation on ice, the mixture was centrifuged at 14,000 × *g* for 10 minutes at 4°C. The supernatant was collected and precipitated with an equal volume of 50% trichloroacetic acid followed by centrifugation at 12,000 × *g* for 20 minutes at 4°C. The histone pellet was washed once with ice-cold acetone containing 0.3 M HCl and once with ice-cold 100% acetone, then dried at 50°C. Dried pellets were resuspended in water supplemented with 20 mM Tris–HCl (pH 8.0), 0.4 N NaOH, and protease inhibitors (7.5 μM aprotinin, 0.5 mM leupeptin, 250 μM bestatin, and 25 mM AEBSF–HCl).

Protein concentration was determined using the Bradford assay (Bio-Rad). Thirty micrograms of acid-extracted histones were prepared in 1× LDS sample buffer (Invitrogen) containing 10% β-mercaptoethanol and separated by 16% sodium dodecyl sulfate–polyacrylamide gel electrophoresis (SDS–PAGE). Proteins were transferred to nitrocellulose membranes and blocked in 1× Tris-buffered saline with Tween-20 (TBST) containing 5% milk for 1 hour. Membranes were incubated overnight at 4°C with primary antibodies against H3K27me3 (Cell Signaling Technology, 9733S; 1:1000) or H3 (Abcam, ab1791; 0.1 μg/mL). The next day, membranes were incubated with the appropriate secondary antibodies for 1 hour at room temperature. Proteins of interest were visualized using the Odyssey imaging system (LI-COR) and quantified using ImageJ.

### Seahorse

Extracellular flux analysis of the Mito Stress Test and Glycolysis Stress Test was performed on the Seahorse XFe96 Analyzer (Seahorse Bioscience) following manufacturer recommendations as described in Lane 2025^63^. Neurons, astrocytes, and radial glia were plated at a density of 100,000 cells/well on Seahorse XF96 V3-PS Microplates (Agilent Technologies, 101085-004). Neurons, astrocytes and radial glia were isolated from fetal cortex (GW18-21) as described above. Cells were plates at a density of 100,000 cells per well. Cells were exposed for 3 days as described above.

The day before the Seahorse experiment, XFe96 extracellular flux assay kit probes (Agilent Technologies, 102416-100) were incubated with the included manufacturer calibration solution overnight at 37°C without CO_2_ injection. The day of the experiment, wells were washed twice in the appropriate Seahorse Media and incubated at 37°C without CO_2_ injection for 1 hour.The Mito Stress Test Media consisted of Seahorse XF base media (Agilent Technologies, 102353-100) with the addition of 2 mM L-glutamine (HyClone, SH30034.01), 1 mM sodium pyruvate (Sigma, S8636), and 10 mM D-glucose (Sigma, G8769), and the Glycolysis Stress Test media consisted of Seahorse XF base media supplemented with 2 mM L-glutamine. Injection ports in the flux plate were loaded with 10-fold concentrated solutions of the drugs corresponding to each assay and calibrated. Seahorse drugs were used at these final concentrations: oligomycin A (2 μM, Sigma 75351-5MG), FCCP (0.75 μM, Sigma C2920), rotenone (0.5 μM, Sigma R8875), antimycin A (0.5 μM, Sigma A8674-25MG), glucose (10 mM), and 2-deoxyglucose (2-DG, 50 mM, Sigma D3179-1G). All drugs were dissolved in DMSO and diluted in the appropriate Seahorse media except for 2-DG, which was prepared in the glycolysis stress test media, stored at -20C, and thawed immediately before use within 1 month of preparation. After calibration, the flux plate containing calibrant solution was exchanged for the Seahorse cell culture plate and equilibrated. The flux analyzer protocol included three basal read cycles and three reads following injection of three distinct combinations of drugs: oligomycin A, FCCP, and rotenone mixed with antimycin A for the Mito Stress Test and D-glucose, oligomycin A, and 2-DG for the Glycolysis Stress Test. Each read cycle included a 3-minute mix cycle followed by a 3-minute read cycle where oxygen consumption rate (OCR) and extracellular acidification rate (ECAR) were determined over time.

In all experiments, OCR and ECAR readings in each well were normalized by protein concentration in the well. Cells were washed twice with phosphate buffered saline (Corning 21-040-CV) supplemented with 1 mM MgCl_2_ and 100 μM CaCl_2_ and lysed in Buffer A, containing 150 mM NaCl, 10 mM HEPES, 1 mM ethylene glycol-bis(β-aminoethylether)-*N*,*N*,*N*′,*N*′-tetraacetic acid (EGTA), and 0.1 mM MgCl2, pH 7.4 with 0.5% Triton X-100 (Sigma, T9284) and Complete anti-protease (Roche, 11245200). Protein concentration was measured using the Pierce BCA Protein Assay Kit (Thermo Fisher Scientific, 23227) according to manufacturer protocol. The BCA assay absorbance was read by a BioTek Synergy HT microplate reader using Gen5 software. For data analysis of OCR and ECAR, the Seahorse Wave Software version 2.2.0.276 was used.

Individual data points represent the average values of a minimum of 5 replicates. For the Mito Stress Test, non-mitochondrial respiration was determined as the lowest OCR following injection of rotenone plus antimycin A; basal respiration was calculated from the OCR just before oligomycin injection minus the non-mitochondrial respiration; and maximal respiration was calculated as the maximum OCR of the three readings following FCCP injection minus non-mitochondrial respiration. For the Glycolysis Stress Test, glycolysis was calculated as the difference in ECAR between the maximum rate measurements before oligomycin injection and the last rate measurement before glucose injection; glycolytic capacity was calculated as the difference in ECAR between the maximum rate measurement after oligomycin injection and the last rate measurement before glucose injection; glycolytic reserve was calculated as the glycolytic capacity minus glycolysis; and non-glycolytic acidification was defined as the last rate measurement prior to glucose injection. The fold-change of each metabolic parameter following PbCl_2_-low treatment, i.e. Pb-treated values normalized to the untreated values for the same cell type for a given experiment, is presented.

## Acknowledgements

We would like to thank the Sloan Laboratory and Dr. Michael Caudle for helpful discussions. This study was supported by the National Institute of Health under grants DSPAN K00 ES033033 (MMS), HERCULES P30 ES019776 (SAS, DBB, PED), NIMH R01 MH125956 (SAS), NIH grant R01ES034796 to EW, 1F31NS127419 (ARL), NIH R01 ES034796 (VF and EW), NIH/NINDS R01NS109025 (YZ), NIH/NIDA R01DA059873 (YZ), NIH/NICHD P50HD103557 (YZ), NIH/NIGMS R35GM150587 (JMS). MMS was supported by Burroughs Wellcome Fund PDEP (MMS). ARL was supported by an ARCS Foundation Award, Robert W. Woodruff Fellowship, and John B. Lyon Memorial Scholarship Award. YZ was also supported by the Broad Stem Cell Research Center Innovation Award (YZ), UCLA Jonsson Comprehensive Cancer Center and BSCRC Ablon Scholars Award (YZ), Rose Hills Foundation Stem Cell Innovation Award (YZ). This study was supported in part by the Emory Integrated Genomics Core (EIGC) and Emory University School of Medicine Flow Cytometry Core which are subsidized by the Emory University School of Medicine and are of the Emory Integrated Core Facilities.

## Contributions

MMS and SAS designed all experiments and wrote the manuscript with input from all contributing authors. MMS performed analysis with support from ANV, EJH and NK. PED performed MS Pb quantification under the supervision of DBB. MMS, AK and AS performed hiPSC cultures, organoid formation and maintenance. MMS and AS performed single-cell captures and library preparation. Xenograft experiments including single-cell captures were performed by LP, RK, and DA under the supervision of YZ and AB. SJS performed live imaging, live-dead, and TUNEL experiments. MMS, SJS, EJH and LP performed Pb exposures. LN, DVF and SNL performed immunoblots under the supervision of SAS and JMS. ARL and EW performed Seahorse assays under the supervision of VF.

## Data Availability

All data underlying data visualizations are provided in “Source_Data” file. Sequencing files (Fastq,.bam, featurecounts, R objects) will be made available on GEO (Accession codes will be available before publication).

## Statistical Tests

Statistical tests were performed in GraphPad Prism (Version 10) or R Studio (Version 4.2.2). All statistical tests and related details (intervals, effect sizes, degrees of freedom and P values) are included in the Source_Data file. We typically employed the Brown-Forsythe test to compare variances across multiple groups. For groups with unequal variances, we used Brown-Forsythe ANOVA or Welch’s ANOVA test. Multiple comparisons were typically performed with Dunnett’s multiple comparisons test to compare experimental groups to control and adjusted P values are reported. For normalized data, we used one sample t test and wilcoxon test to compare experimental groups to a hypothetical control. All reported P values are two-tailed.

## Code Availability

R code for analysis and figure generation will be made available on the Sloan Lab github before publication (https://github.com/sloanlab-emory).

---

**Extended Data Fig. 1.**
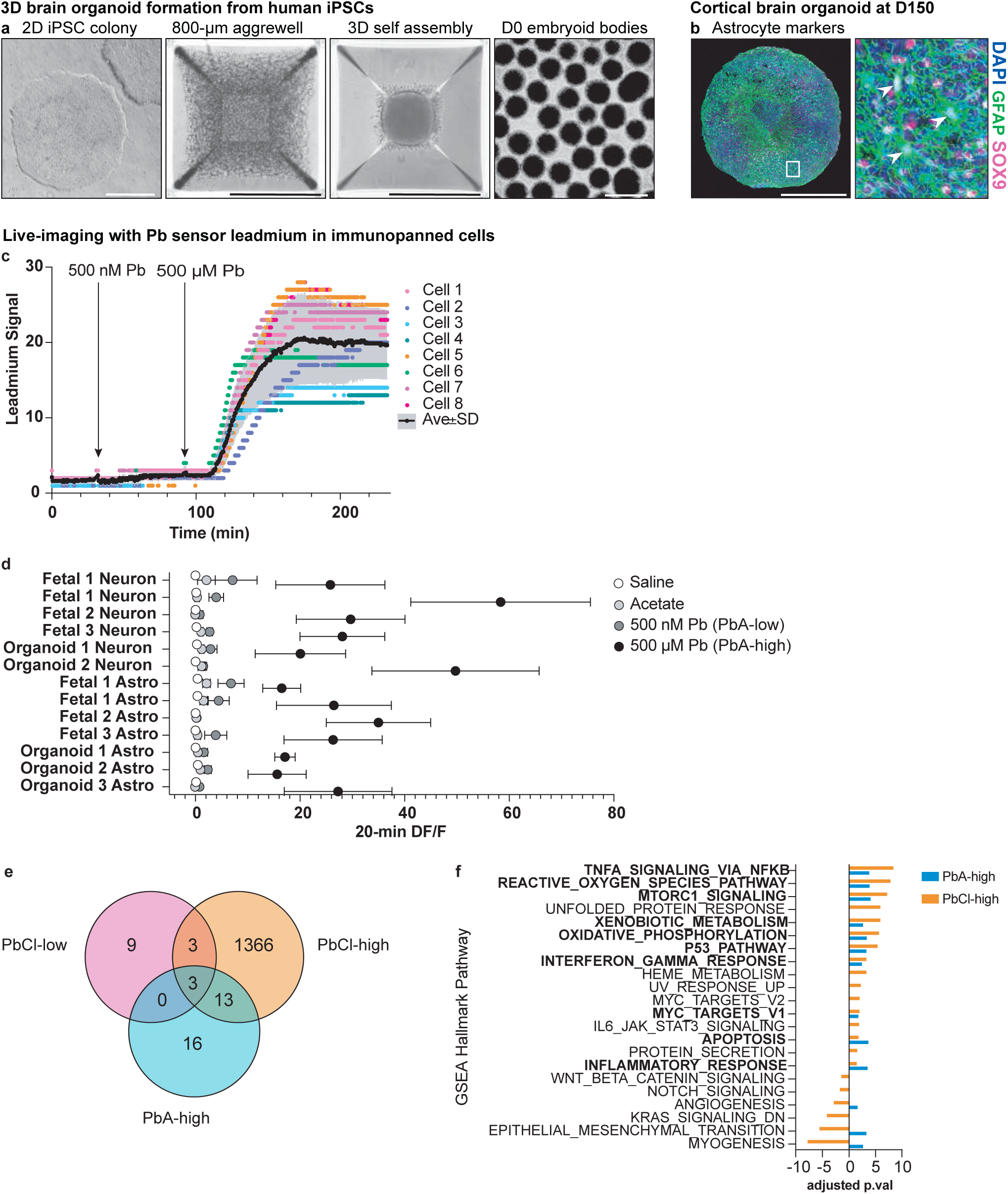
**a,** Organoids are formed from human induced pluripotent stem cells (hiPSCs) cells that self-assemble into 3D spheres after 24 hours in 800 µm aggrewells. **b,** As they mature, cortical organoids spontaneously generate astrocytes (green GFAP, magenta SOX9) as shown at D150. Scale bars in a-b are 500 µm. c, Leadmium recordings for 8 astrocytes isolated from a GW18 fetal cortex, **d,** 20-min DF/F values are shown for control additions of saline, acetate, 500 nM PbA and 500 µM PbA. **e,** Venn diagram showing whole organoid transcriptional response to PbA and PbCh high and low doses, **f,** GSEA hallmark pathways for PbA-high (500 µM) and PbCh-high (500 µM).

**Extended Data Fig. 2.**
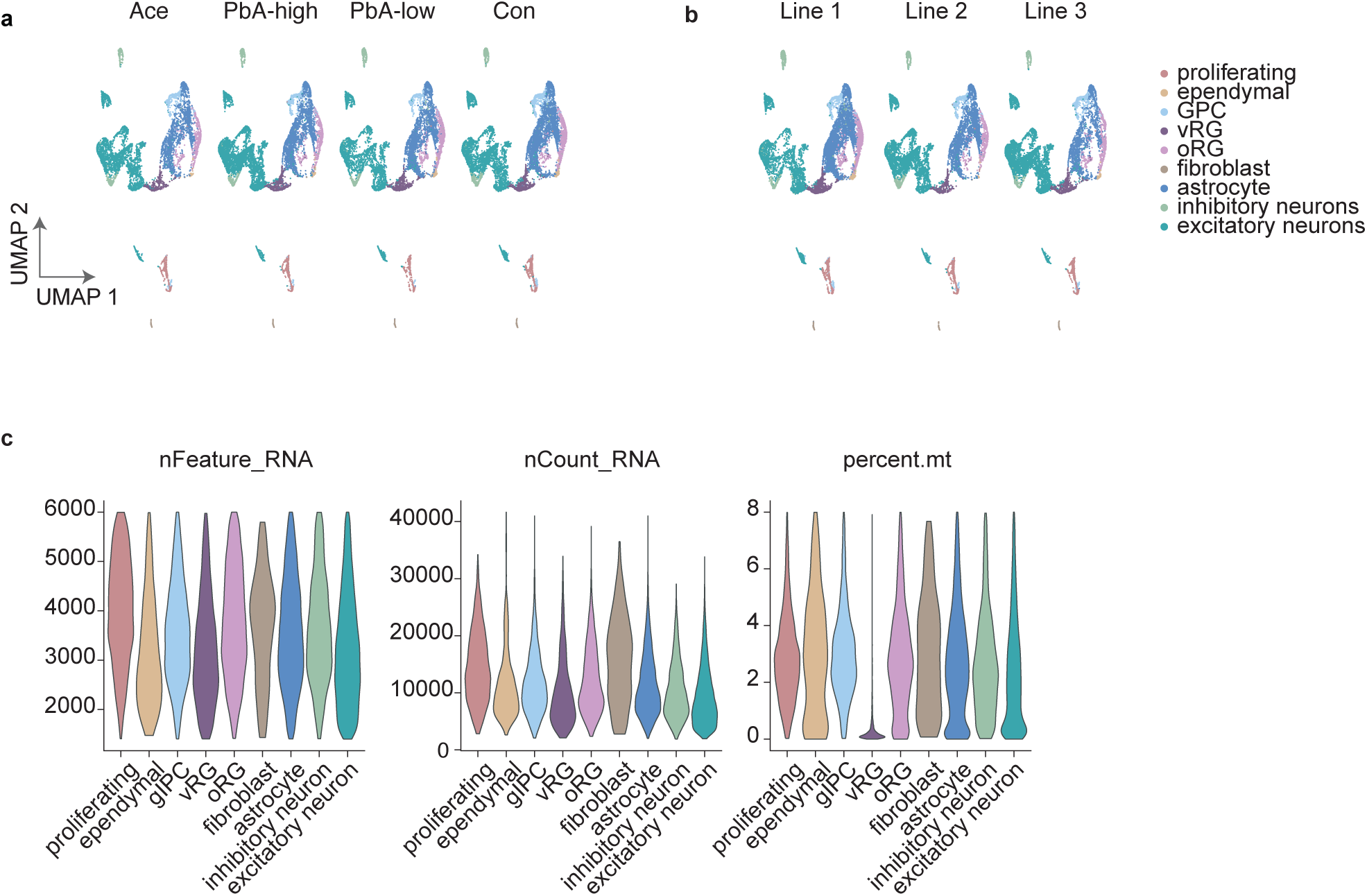
**a,** Single-cell data visualized on a UMAP split into each exposure group: Acetate, Pb-high (500 µM), Pb-low (500 nM), and Control (normal media), b, Single-cell data visualized on a UMAP split by organoid hiPSC line: Line 1, Line 2, Line 3. c, Quality control metrics by cell cluster: nFeature, nCount, percent mitochondrial reads.

**Extended Data Fig. 3.**
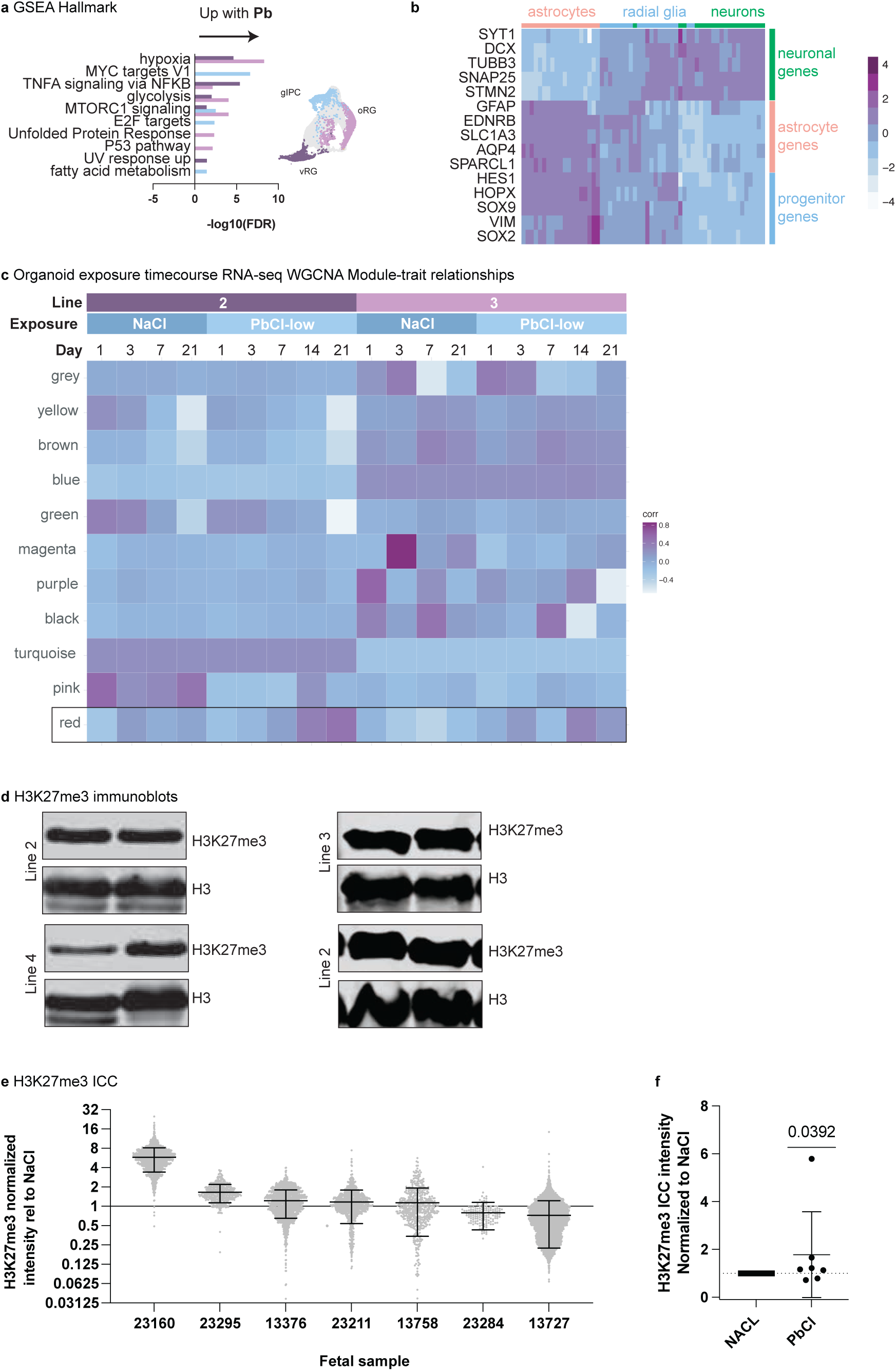
**a,** GSEA Hallmark pathways altered by Pb in progenitors (ventral radial glia, glial progenitor cells, outer radial glia), b, RNA-Seq from immunopanned cells showed high expression of neuronal genes in neurons and astrocyte genes in astrocytes. Radial glia expressed markers of neural progenitors as well as lower levels of astrocyte and neuron genes, c, WGCNA module-trait reíationships for PTPRZ1 immunopanned cells, **d,** H3K27me3 immunoblot images for data shown in Fig 3g. **e,** H3K27me3/H3 immunoblot quantification for PbCI2 high.

**Extended Data Fig. 4.**
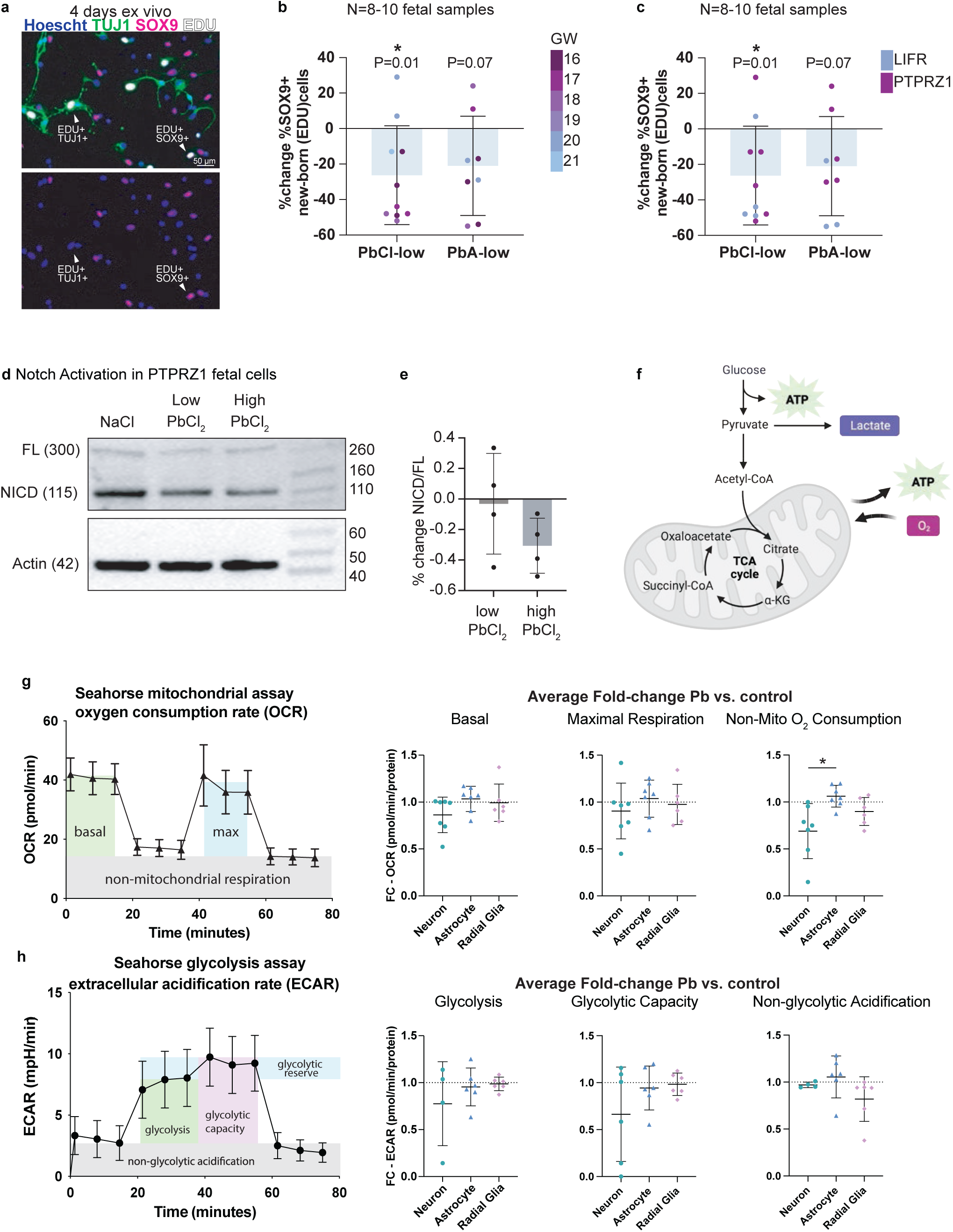
**a**, Fetal radial glia stained for TUJ1 (green) *ex vivo*, SOX9 (magenta) and EdU (white). **b**, Gestational week metadata for individual fetal samples in Fig 4F, percent change in the %SOX9+ cells for N=8-10 individual fetal samples. **c**, Immunopanning antibody metadata for individual fetal samples in Fig 4F, percent change in the %SOX9+ cells for N=8-10 individual fetal samples. **d**, Representative immunoblot for full length NOTCH (FL), NOTCH intracellular domain (NICD), and actin. **e**, In N=4 fetal samples, the precent change in the NICD/FL ratio, and indicator of NOTCH activity. **f-h**, Bioenergetic processes measured in the Seahorse extracellular flux oximetry assays are oxygen consumption and extracellular acidification rate, which is largely attributed to lactate accumulation. Seahorse assays were performed in fetal neurons, astrocytes, and radial glia 4 days ex vivo treated with PbCl_2_-low for 3 days. **g**, The Seahorse Mito Stress Test is used to calculate mitochondrial respiratory parameters based on oxygen consumption rate (OCR). FC-OCR is shown for basal respiration, maximal respiration, and non-mitochondrial respiration (N=6-7). For non-mitochondrial respiration, P=0.0165 by Brown-Forsythe ANOVA test and P=0.039 Dunnett’s T3 multiple comparisons test for astrocytes vs. neurons. **h**, The Seahorse Glycolysis Stress Test is used to calculate glycolytic parameters based on extracellular acidification rate (ECAR). FC-ECAR is shown for glycolysis, glycolytic capacity, and glycolytic reserve (N=2-4). All data are presented as average fold-change of Pb-treated cells to untreated -treated cells ± SD (see Methods). for PTPRZ1 immunopanned cells, **d,** H3K27me3 immunoblot images for data shown in Fig 3g. **e,** H3K27me3/H3 immunoblot quantification for PbCI2 high.

**Extended Data Fig. 5.**
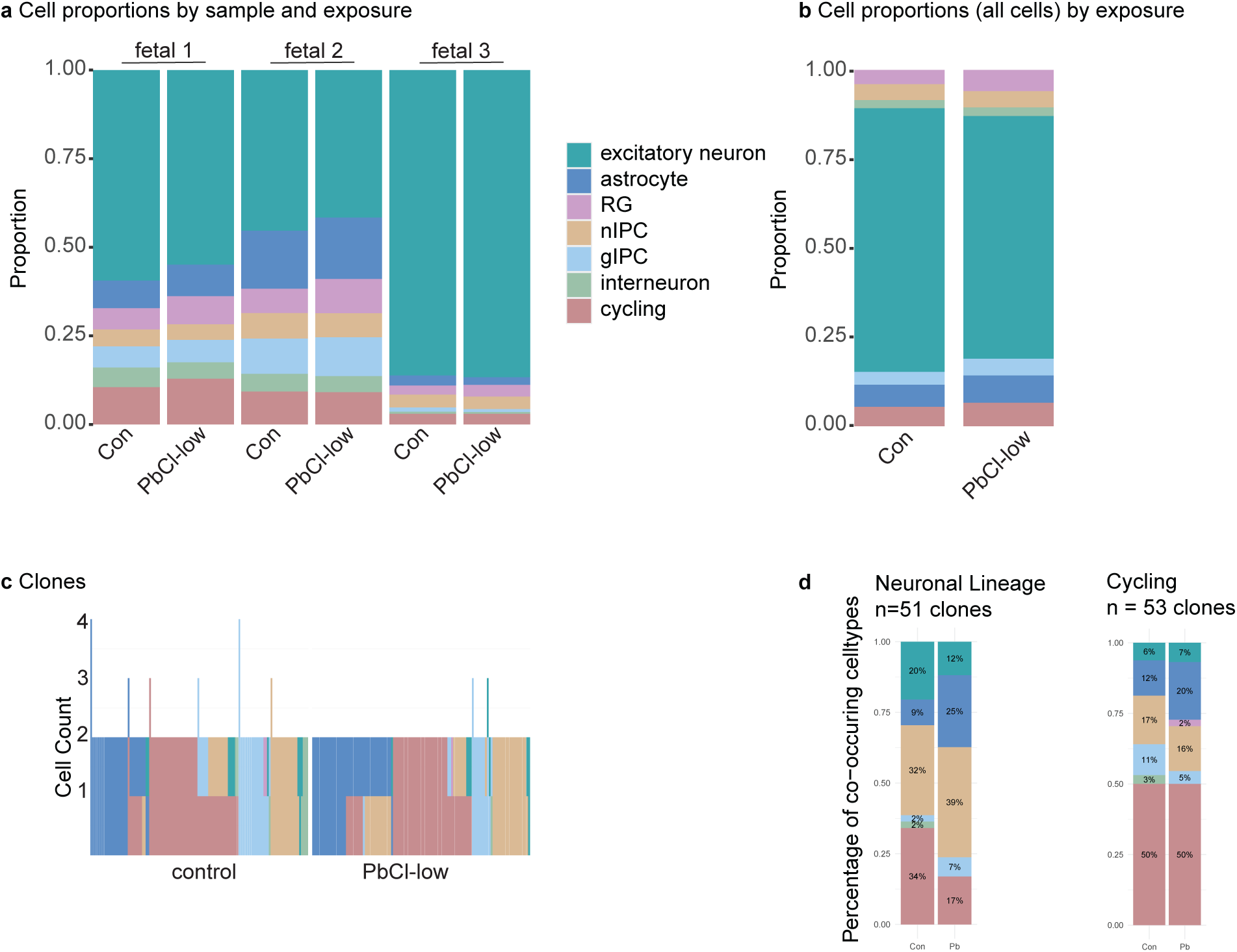
**a-b** Cell proportions for CellTag data set separated by sample and exposure, **c,** Individual clones for each condition, d, The composition of mixed clones containing nIPCs (left) or cycling cells (right), both normalized by the total cells in the clone group.

**Extended Data Fig. 6.**
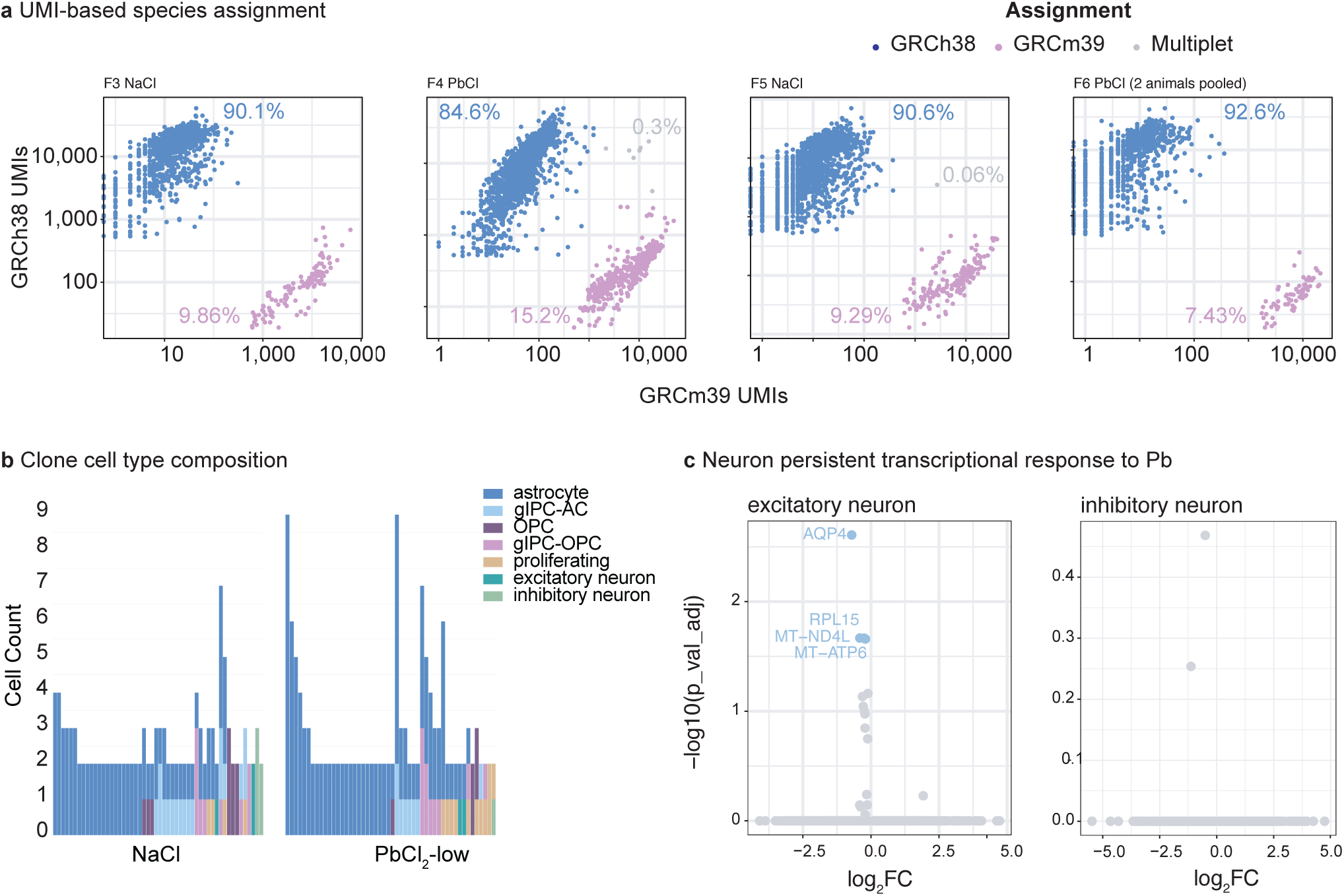
**a** single-cell species assignment visualized by GRCh38 (human) vs GRCm39 (mouse) UMIs b, Individual clones for each condition, c, Neuron transcriptional response to Pb.

